# Privacy-Preserving Federated Neural Network Learning for Disease-Associated Cell Classification

**DOI:** 10.1101/2022.01.10.475610

**Authors:** Sinem Sav, Jean-Philippe Bossuat, Juan R. Troncoso-Pastoriza, Manfred Claassen, Jean-Pierre Hubaux

## Abstract

Training accurate and robust machine learning models requires a large amount of data that is usually scattered across data-silos. Sharing or centralizing the data of different healthcare institutions is, however, unfeasible or prohibitively difficult due to privacy regulations. In this work, we address this problem by using a novel privacy-preserving federated learning-based approach, *PriCell*, for complex machine learning models such as convolutional neural networks. *PriCell* relies on multiparty homomorphic encryption and enables the collaborative training of encrypted neural networks with multiple healthcare institutions. We preserve the confidentiality of each institutions’ input data, of any intermediate values, and of the trained model parameters. We efficiently replicate the training of a published state-of-the-art convolutional neural network architecture in a decentralized and privacy-preserving manner. Our solution achieves an accuracy comparable to the one obtained with the centralized solution, with an improvement of at least one-order-of-magnitude in execution time with respect to prior secure solutions. Our work guarantees patient privacy and ensures data utility for efficient multi-center studies involving complex healthcare data.

## Introduction

Machine learning models, in particular neural networks, extract valuable insights from data and have achieved unprecedented predictive performance in the healthcare domain, e.g., in single-cell analysis^1^, aiding medical diagnosis and treatment^2,3^, or in personalized medicine^4^. Training accurate and unbiased models without overfitting requires access to a large amount of diverse data that is usually isolated and scattered across different healthcare institutions^5^. Sharing or transferring personal healthcare data is, however, often unfeasible or limited due to privacy regulations such as GDPR^6^ or HIPAA^7^. Consequently, privacy-preserving collaborative learning solutions play a vital role for researchers, as they enable medical advances without the information about each institution’s data being shared or leaked. Collaborative learning solutions play a particularly important role for studies that involve novel informative, yet not universally established, data modalities such as high dimensional single-cell measurements, where the number of examples is typically low at individual study centers and only amounts to critical mass for the successful training of machine learning models across multiple study centers^8^. The ability to satisfy privacy regulations in an efficient and effective manner constitutes a pivotal requirement to carry out translational multi-center studies.

Federated learning (FL) has emerged as a promising distributed learning approach, where the parties keep their raw data on their premises and exchange intermediate model parameters^9,10^. This approach has enabled collaborative learning for several medical applications, and it has been shown that FL performs comparably to centralized training on medical datasets^11–13^. Recently, the concept of swarm learning (SL) has been proposed; it enables decentralized machine learning for precision medicine. The seminal work of SL^14^ is based on edge computing and permissioned blockchains and removes the need for a central server in the FL approach. Despite the advantages of FL and SL for keeping the sensitive data local and for reducing the amount of data transferred/outsourced, the model and the intermediate values exchanged between the parties remain prone to several privacy attacks executed by the other parties or the aggregator (in FL), such as membership inference attacks^15,16^ or reconstructing the parties’ inputs^17–19^. In this work, we provide a solution that further conceals the global machine learning model from the participants by relying on mathematically secure cryptographic techniques to mitigate these inference attacks.

In order to mitigate or prevent the leakage in the FL setting, several privacy-preserving mechanisms have been proposed. These mechanisms can be classified under three main categories, depending on the strategy they are based on: differential privacy (DP), secure multiparty computation (SMC), and homomorphic encryption (HE).

Differential privacy (DP)-based solutions aim to perturb the parties’ input data or the intermediate model values exchanged throughout the learning. Several studies in the medical domain keep the data on the local premises and use FL with a differential privacy-mechanism on the exchanged model parameters^20–22^. Despite being a pioneering mitigation against privacy attacks, DP-based solutions perturb the model parameters, thus decreasing the utility and making the deployment harder for medical applications, where the accuracy is already constrained by limited data. Quantification of the privacy achieved via DP-based approaches is also very difficult^23^ and the implementation of DP, especially in medical imaging applications, is not a trivial task^5^.

Another line of research relies on secure multiparty computation (SMC) techniques to ensure privacy and to enable collaborative training of machine learning models^24–28^. SMC techniques rely on secret-sharing the data of the parties and on performing the training on the secret-shared data among multiple computing nodes (usually 2,3, or 4 nodes). Nevertheless, it is usually hard to deploy these solutions, as they often rely on a trusted third party for the sake of efficiency. Moreover, their scalability with the number of parties is poor due to the large communication overhead.

Finally, several works employ homomorphic encryption (HE) to enable secure aggregation or to secure outsourcing of the training to a cloud server^29,30^. These solutions, however, cannot solve the distributed scenario where parties keep their local data in their premises.

The adoption of each of the aforementioned solutions introduces several privacy, utility, and performance trade-offs that need to be carefully balanced for healthcare applications. To balance these trade-offs, several works employ multiparty homomorphic encryption (MHE)^31,32^. Although the underlying model in these solutions enables privacy-preserving distributed computations and maintains the local data of the parties on their local premises, the functionality of these works is limited to the execution of simple operations, i.e., basic statistics, counting, or linear regression and the underlying protocols do not support an efficient execution of neural networks in the federated learning setting.

Recently, Sav et al. proposed a more versatile solution, POSEIDON, for enabling privacy-preserving federated learning for neural networks by relying on MHE^33^ to mitigate FL inference attacks by keeping the model and intermediate values encrypted. Their solution, however, does not address the implementation and efficient execution of convolutional neural networks, a widely adopted machine learning model to analyze complex data types such as single-cell data.

We propose *PriCell*, a novel solution based on MHE to enable the training of a federated convolutional neural network in a privacy-preserving manner, thus preserving the utility of the data for single-cell analysis. To the best of our knowledge, *PriCell* is the first of its kind in the regime of privacy-preserving multi-center single-cell studies under encryption. By bringing privacy-by-design and by preventing the transfer of patients’ data to other institutions, our work contributes to single-cell studies and streamlines the slow and demanding process of the reviewing of independent ethics committees for consent forms and study protocols. To mitigate FL attacks, we keep the model and any value that is exchanged between the parties in an encrypted form, and we rely on the threat model and setting proposed in the work of Sav et al.^33^ (detailed in the Methods section).

By designing new *packing* strategies and homomorphic matrix operations, we improve the performance of the protocols for encrypted convolutional neural networks (CNNs) that are predominantly used in the healthcare domain^34^. To evaluate our system within the framework of single-cell analysis, we train a convolutional neural network (CellCnn), designed by Arvaniti and Claassen^35^, within our privacy-preserving system for the disease classification task. We also show the feasibility of our solution with several single-cell datasets utilized for cytomegalovirus infection (CMV)^36^ and acute myeloid leukaemia (AML)^37^ classification, and one dataset for non-inflammatory neurological disease (NIND) and relapsing–remitting multiple sclerosis (RRMS)^38^ classification. We compare our classification accuracy in a privacy-preserving federated learning setting with the centralized and non-encrypted baseline. Our solution converges comparably to the training with centralized data, and we improve on the state-of-the-art decentralized secure solution^33^ in terms of training time. For example, in a setting with 10 parties, we improve POSEIDON’s execution time by at least one order of magnitude.

## Results

In this section, we introduce the system overview of our solution, present the neural network architecture that is used for our evaluation, and lay out our experimental findings.

### System Overview

We summarize *PriCell*’s system and its workflow for collaborative training and query evaluation (prediction), in Figure 1a. Assuming there are four healthcare institutions with each holding its respective *secret key*, the workflow starts with the generation of a *collective public key* and a set of *evaluation keys* that are necessary for the encrypted operations, using each participant’s secret key. We refer to this phase as the *setup phase*. In the second phase, the participants agree on the initial random global model weights (*W*_*g*_) and encrypt them with the collective public key. We denote the encryption of any value with boldface letters, i.e., ***W*** _***g***_. After encrypting the initial global weights, the *local computation phase* begins; we base this phase on a variant of the FedAvg algorithm^9^ to enable collective training. Based on this algorithm, to find the model gradients (**∇*W*** _***k***_), each party performs several encrypted training iterations on their local data (local iteration). The local-model gradients are then sent and aggregated at one of the parties that will perform the global model update. The updated model is then broadcast back and the process returns to Phase 2. After a fixed number of training iterations, the participants can choose to keep the model confidential (option 5.1 in Figure 1a) or to decrypt it for further analysis (option 5.2 in Figure 1a). If prediction-as-a-service is offered to a querier (a researcher) and the model is kept encrypted, the querier must encrypt the evaluation data (***X***_***q***_) with the collective public key of the parties. Once the prediction is done, the result (**ŷ**) is collectively switched to the public key of the querier by using the underlying cryptoscheme’s *collective key switching* functionality. If the model is instead decrypted, the querier encrypts the data with their own key hence no key switch is needed after the prediction. As a result, regardless of the model being confidential or not, the evaluation data of the querier and the prediction result always remain protected, as only the querier can decrypt the end result.

**Figure 1.**
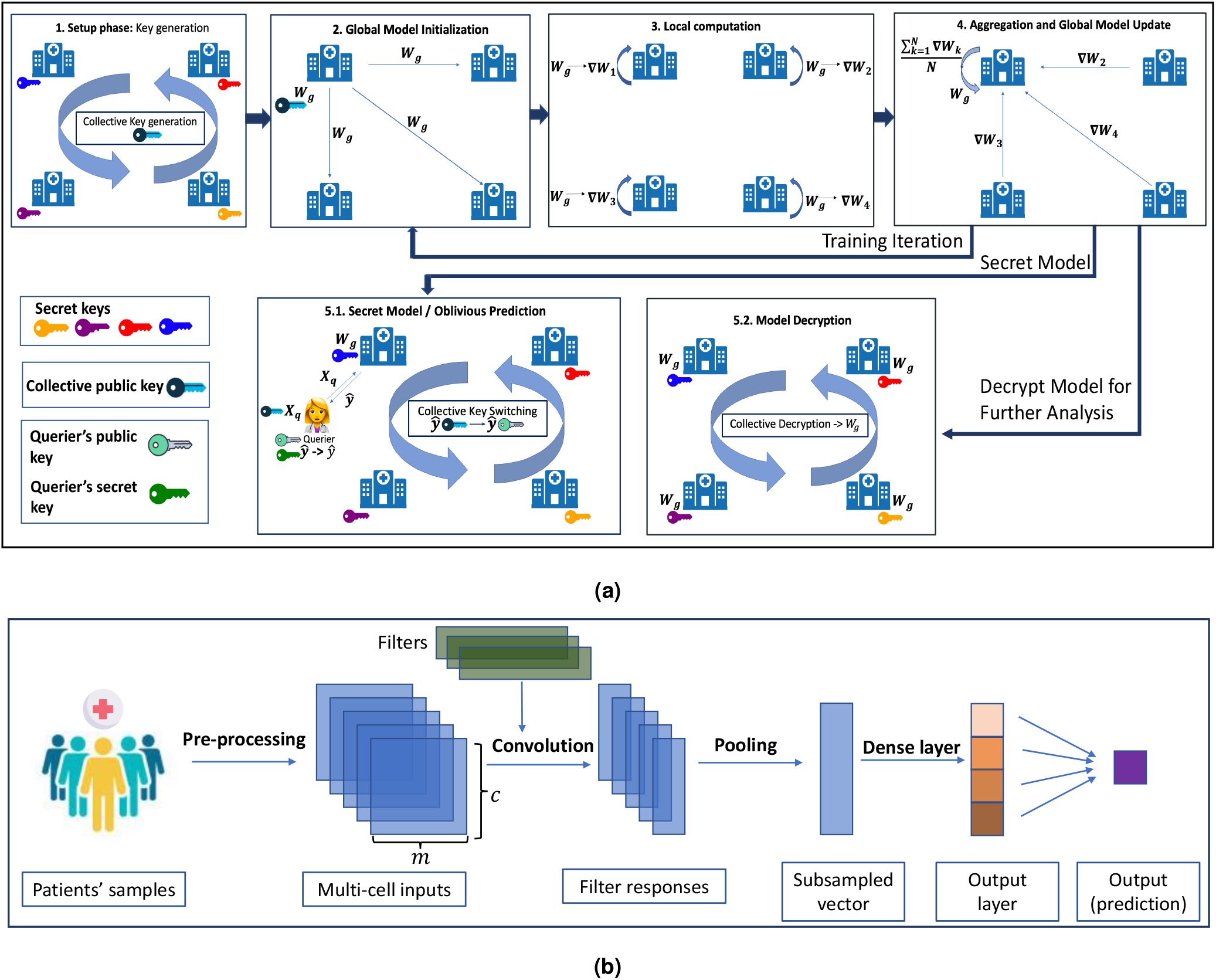
**(a)** *PriCell*’s Training and Evaluation Workflow. Training encapsulates the generation of cryptographic keys and the federated learning iterations on an encrypted model with multiple healthcare institutions. After training, the model is either kept encrypted or decrypted for further analysis. **(b)** CellCnn^35^ Neural Network Architecture that is used in local computation phase of **(a)**. The network takes multi-cell samples as an input and applies a 1D convolution with *h* filters followed by a pooling layer. A dense (fully-connected) layer then outputs the phenotype prediction.

### CellCnn Model Overview

CellCnn is a convolutional neural network that enables multi-instance learning and associates a set of observations on cellular population, namely multi-cell inputs, with a phenotype^35^. This architecture is designed for detecting rare cell subsets associated with a disease, by using multi-cell inputs generated from high-dimensional single-cell marker measurements. By their nature, these inputs can be used to predict the phenotype of a donor or the associated phenotype for a given subset of cells. In this scenario, we enable privacy-preserving and distributed multi-instance learning, and we compare our classification performance with the baseline (CellCnn^35^ trained on centralized data with no privacy protection). We note here that replicating the full-pipeline of CellCnn^35^ for downstream analysis requires either heavy approximations under encryption or the decryption of the trained model. Our solution enables the collective and privacy-preserving training for the classification task, whereas subsequent analyses that require access to the model are out-of-the-scope of this work. Yet, we show the negligible effect that our encryption would practically have on these analyses in Supplementary Note 6.

We show the architecture of CellCnn^35^ in Figure 1b. The network comprises a 1D convolutional layer followed by a pooling layer and a dense (fully-connected) layer. Each multi-cell input sample in Figure 1b is generated using *c* cells per phenotype with *m* features (markers), and these samples are batched to construct multi-cell inputs. The training set is then generated by choosing *z* multi-cell inputs per output label or per patient.

We refer the reader to the work of Arvaniti and Claassen^35^ for the details of the neural network architecture. We detail the changes we introduce to this architecture to enable operations under homomorphic encryption in the Local Neural Network Operations subsection of the Methods.

### Experimental Evaluation

We evaluate our proposed solution in terms of model accuracy, runtime performance, scalability with the number of parties, number of data samples, number of features, and communication overhead. And we provide a comparison with the prior work in this section. We give details on the machine learning hyperparameters and security parameters used for our evaluation in the Supplementary Note 3.

#### Model Accuracy

To assess our solution in terms of accuracy, we use the same datasets used in two peer-reviewed biomedical studies^35,38^. We rely on three datasets to perform non-inflammatory neurological disease (NIND), relapsing–remitting multiple sclerosis (RRMS), Cytomegalovirus Infection (CMV), and Acute Myeloid Leukaemia (AML) classification. We give the details of each dataset in the Datasets subsection of the Methods. **Our aim is to show that *PriCell* achieves a classification performance on par with the centralized non-private baseline**.

As the original studies rely on centralized datasets, we evenly distribute the individual donors in the respective dataset over *N* parties. We give the classification performance on these datasets in Figure 2, 3, and 4. The x-axis shows different training approaches: (i) the data is centralized and the original CellCnn approach^35^ is used for the training and classification to construct a baseline, (ii) each party trains a model only with its local data (Local), without collaborating with other parties, and (iii) our solution for privacy-preserving collaboration between parties is used (*PriCell*). For the Local training (ii), we average the test accuracy achieved by individual parties.

**Figure 2.**
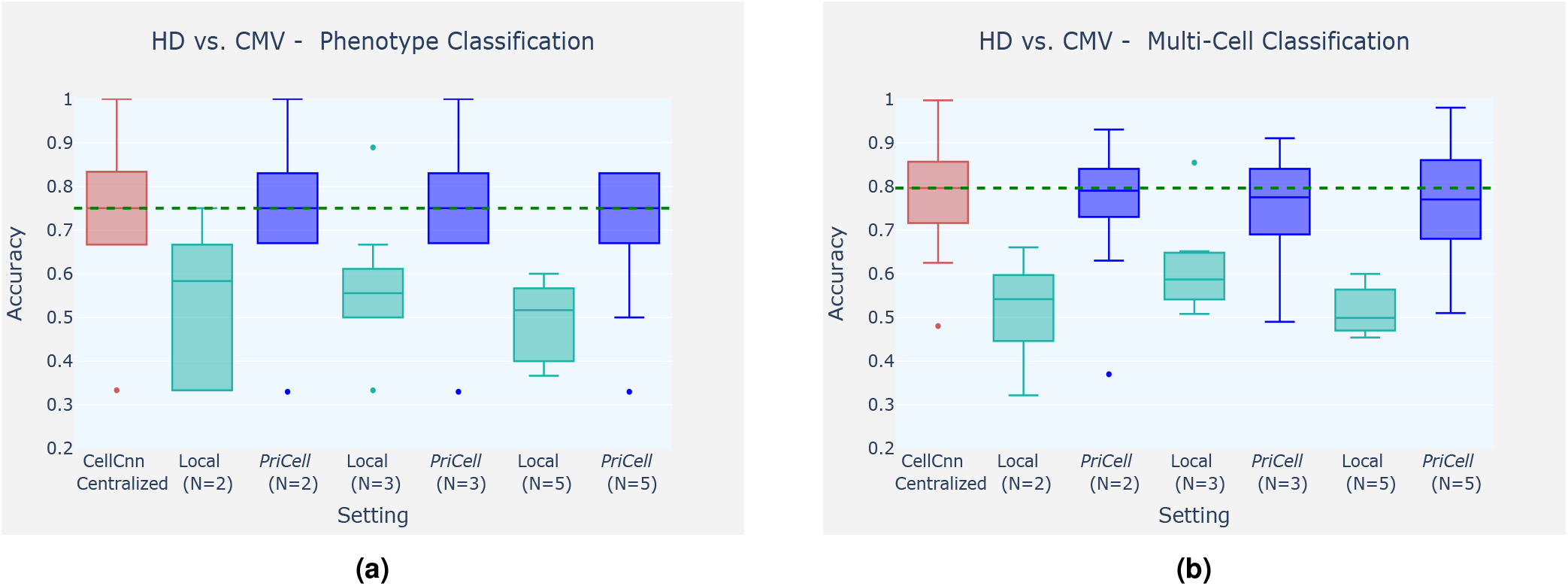
Accuracy Boxplots when classifying healthy donor (HD) vs. cytomegalovirus infection (CMV) for training multi-cells drawn from the bag of all cells per class. Experiments are repeated 10 times with different train and test set splits, the vertical dashed line illustrates the median for the baseline (CellCnn) and the dots represent the outliers. Classification accuracy is reported for two datasets: **(a)** phenotype classification of 6 patients and **(b)** multi-cell input classification on 4000 samples.

**Figure 3.**
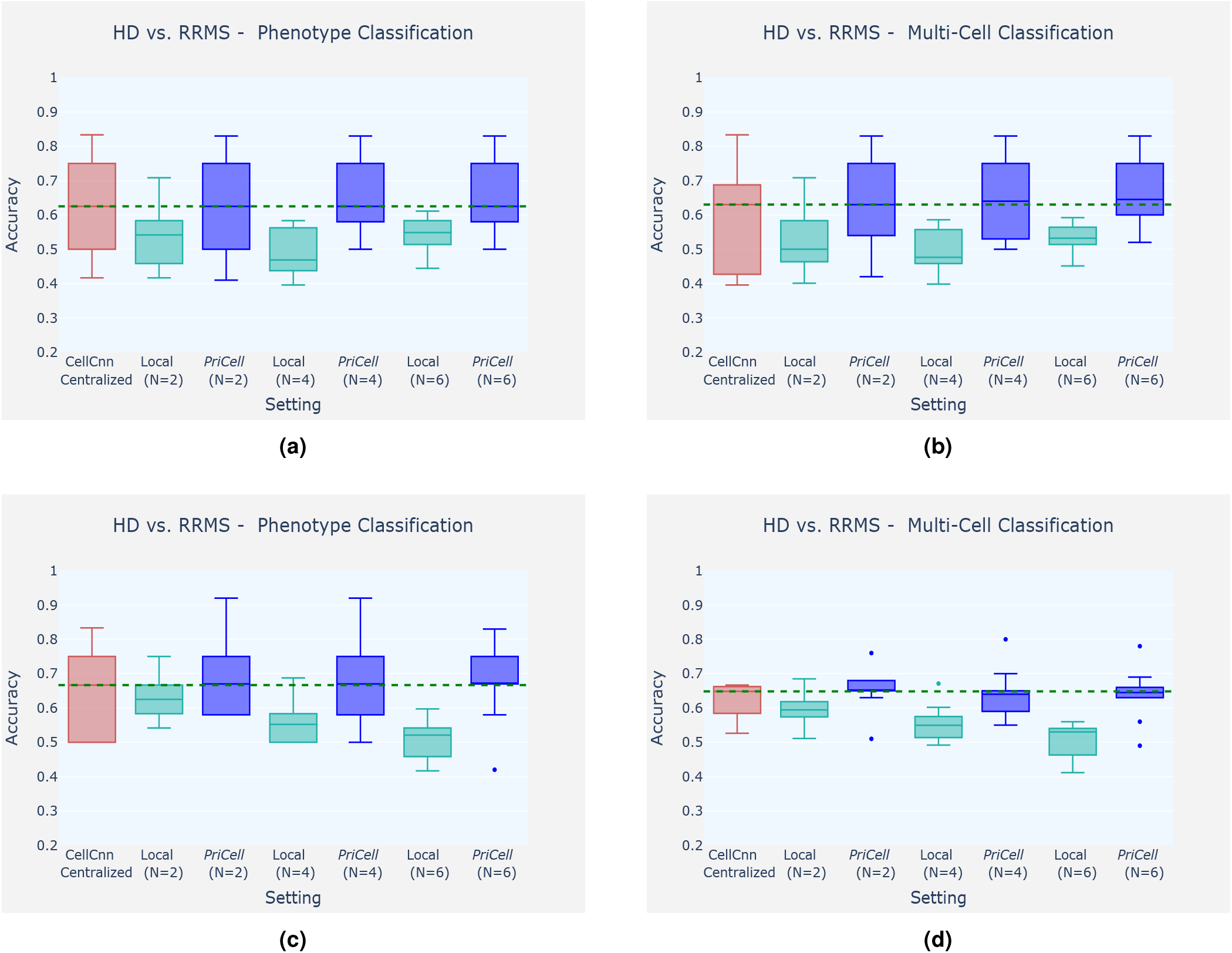
Accuracy boxplots when classifying healthy donor (HD) vs. relapsing–remitting multiple sclerosis (RRMS), for training multi-cells drawn from the bag of all cells per class **(a-b)** and drawn from each patient separately **(c-d)**. Experiments are repeated 10 times with different train and test set splits, the vertical dashed line illustrates the median for the baseline (CellCnn) and the dots represent the outliers. Classification accuracy is reported for two datasets: multi-cell input classification on 96 samples, and phenotype classification of 12 patients.

**Figure 4.**
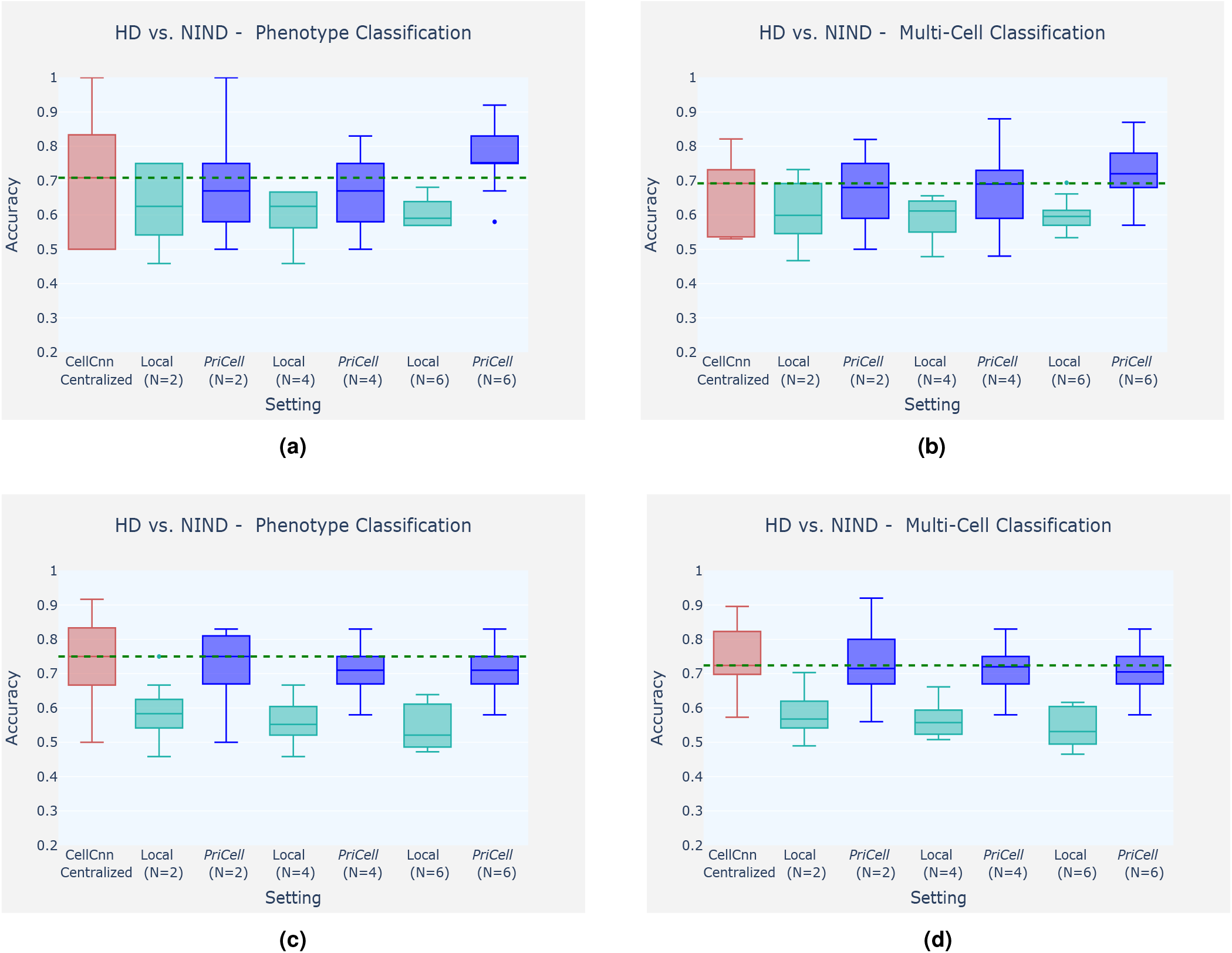
Accuracy Boxplots when classifying healthy donor (HD) vs. non-inflammatory neurological disease (NIND), for training multi-cells drawn from the bag of all cells per class **(a-b)** and drawn from each patient separately **(c-d)**. Experiments are repeated 10 times with different train and test set splits, the vertical dashed line illustrates the median for the baseline (CellCnn) and the dots represent the outliers. Classification accuracy is reported for two datasets: multi-cell input classification on 96 samples and phenotype classification of 12 patients.

In our experiments, random multi-cell inputs that are used for training are drawn with replacement from the original training samples. Drawing multi-cell inputs can be done in two ways: using the bag of all cells per class or individually drawing them from each patient. We report the classification performance by using two test datasets: One set is generated by using multi-cell inputs with *c* = 100 −200 cells drawn from all patients in the test set to increase the size of the test set for multi-cell classification; and the second set is generated by drawing 1000 ™10000 cells from each *donor* separately for phenotype prediction. We give more details about the setting and hyperparameters for each experiment in the Supplementary Note 3.

For CMV classification, we generate the training data by drawing random cell subsets from the cell bags *per phenotype*. For NIND and RRMS classification, we observe that drawing multi-cells per phenotype varies the accuracy between runs and that the median accuracy over 10 runs increases when distributing the initial dataset among *N* = 6 parties (see Figure 3a, 3b and 4a, 4b). This suggests that separately drawing multi-cell inputs from each individual performs better for this task, as corroborated by the results obtained with drawing 2000 cells from *each patient* with replacement (see Figure 3c, 3d and 4c, 4d). Finally, in Supplementary Table **2**, we report the median accuracy, precision, recall, and F-score of 10 runs (with different train and test set splits) on patient-based sub-sampling for NIND and RRMS, and phenotype-based sub-sampling for CMV.

To construct a realistic overall distribution, we limit the number of parties to be lower than the number of donors in the dataset. We observe that, given a sufficient number of samples per party, our distributed secure-solution achieves classification performance comparable to the original work, where the data is centralized and the training is done without privacy-protection. In the experiments on CMV, for example, the median accuracy achieved by *PriCell* is exactly the same as the centralized baseline for phenotype classification and very close (at most 2% gap) for multi-cell classification. Analogous results are obtained for the other experiments: Our privacy-preserving distributed solution achieves almost the same median accuracy with the baseline in RRMS and NIND with patient-based sub-sampling, where the datasets are sufficiently large to be distributed among up to 6 parties.

Lastly, we provide the classification performance on AML in Supplementary Table **2**. As the dataset is relatively small, emulating a distributed setting with more than two parties was not feasible for this task and, as the accuracy does not vary in between different train-test splits, we do not provide the boxplots on the accuracy. However, we observe that with two parties in *PriCell* training, the accuracy remains exactly same as the centralized baseline for AML classification.

Most importantly, our evaluation shows that there is always a significant gain in classification performance when switching from local training to privacy-preserving collaboration. The number of donors that each institution has is insufficient for individually training a robust model. In all experimental settings, for a fixed number of *N, PriCell* achieves better performance than the local training while ensuring the confidentiality of the local data.

#### Runtime

We report in Table 1, the execution times for the training and prediction with *N* = 10 parties and a *ring degree 𝒩* = 2^15^. To be able to compare the runtimes at a larger scale, we use synthetically generated data for this set of experiments and vary the number of features (*m*). We generate a data set of 1000 samples per party with *c* = 200 cells per sample. We use *h* = 8 filters, a local batch size of *n* = 100, and 20 global epochs for training. We report the execution time of the setup phase, of the local computations, and of its communication. We include the execution time of *distributed bootstrapping* (see Methods Section for details) as part of the communication time, which takes 1.2s per iteration and 122s over 20 epochs. Hence, the communication column for training comprises the time to perform all communication between parties throughout the training, distributed bootstrapping, and the model update.

**Table 1.** *PriCell*’s execution times for training and prediction with a varying number of features (*m*), 10 parties, and ring degree *𝒩* = 2^15^ (2^14^ *ciphertext slots*). The computation is single-threaded in a virtual network with an average network delay of 0.17 ms and 1 Gbps bandwidth on 10 Linux servers with Intel Xeon E5-2680 v3 CPUs running at 2.5 GHz with 24 threads and 12 cores and 256 GB RAM.

| $m$ | Training Execution Time [sec] | | | Prediction Execution Time [sec] | | |
| --- | --- | --- | --- | --- | --- | --- |
|  | Setup | Local Computation | Communication | Local computation<br>Querier + Server | Communication | Collective Key-Switch |
| 8 | 17.8 | 753.4 | 370.1 | 0.2 + 0.1 | 0.3 | 0.3 |
| 16 | 18.1 | 778.7 | 387.0 | 0.3 + 0.2 | 0.6 | 0.3 |
| 32 | 19.3 | 836.1 | 393.7 | 0.3 + 0.4 | 1.0 | 0.3 |
| 64 | 21.9 | 951.1 | 373.1 | 0.6 + 0.6 | 2.2 | 0.3 |
| 128 | 24.5 | 1135.9 | 374.8 | 1.6 + 1.5 | 4.2 | 0.3 |

We observe that *PriCell* trains, in less than 20 minutes, a CellCnn model on a training set of 200 cells per sample, 1000 samples per party, and 32 features across 10 parties, including the setup phase and communication. The training time, when the number of features varies, remains 20-25 minutes, which is the result of our efficient use of the SIMD operations provided by the cryptosystem; this is further discussed in the scalability analysis.

We also report in Table 1, the execution times of an oblivious prediction when both the model and the data are encrypted (Phase 5.1 of Figure 1a). We recall that the collective key-switching operation enables us to change the encryption key of a ciphertext from the parties’ collective key to the querier’s key. The maximum number *n* of samples that can be batched together for a given ring degree *𝒩*, number of labels *o*, and number of features *m*, is (*𝒩 /*2)*/*(*m* · *o*) (we also need *m/*2 ciphertexts to batch those samples, see Supplementary Note 4 for more details). Hence, in our case the maximum prediction batch size for *𝒩* = 2^15^ and *o* = 2 is *n* = 2^13^*/m*.

We observe that the local computation for the prediction increases linearly with *m*, and is linked to the cost of the dominant operation, the convolution, which is, unlike training, carried out between two encrypted matrices (see Supplementary Note 4). The communication required for prediction includes *m/*2 ciphertexts sent by the querier and one ciphertext (prediction result) sent back by the server. Hence, the communication time also increases linearly with *m*. Lastly, the time for the collective key-switch remains constant, as it is performed once at the end of the prediction protocol on only one ciphertext.

#### Scalability Analysis

Figure 5 shows the scalability of *PriCell* with the number of parties, the global number of rows (samples), number of features (markers), and the number of filters for one global training epoch that is to process once all the data of all parties. Unless otherwise stated, we use *c* = 200 cells per sample, a local batch size of *n* = 100, *m* = 38 features, and *h* = 8 filters, for all settings. We first report the runtime with an increasing number of parties (*N*) in Figures 5a and 5b when the global number of data samples is fixed to *s* = 18000 and when the number of samples per party is fixed to 500, respectively. As the parties perform local computations in parallel, *PriCell*’s runtime decreases with increasing *N* when *s* is fixed (Figure 5a).

**Figure 5.**
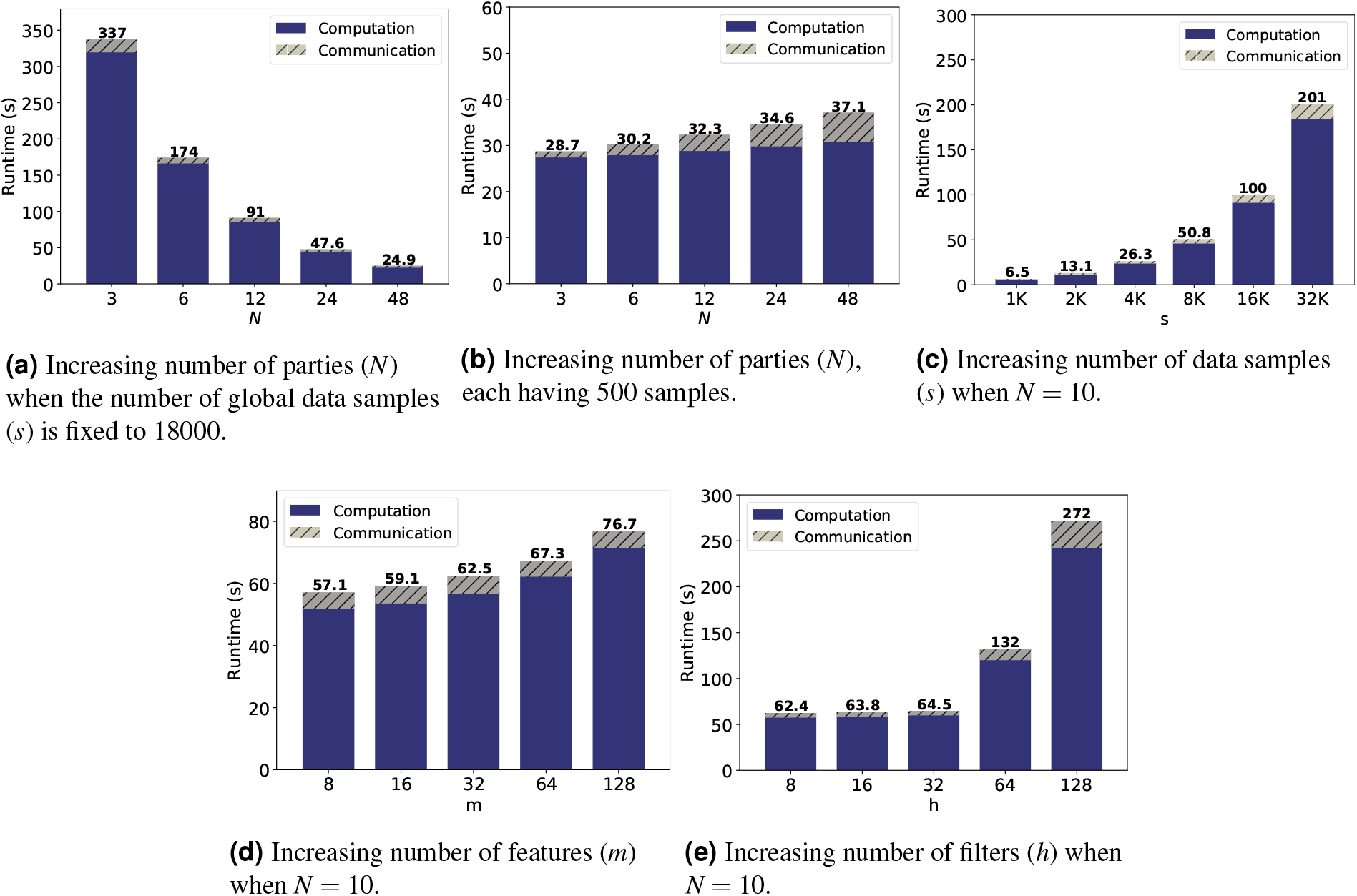
*PriCell*’s training execution time and communication overhead for one training epoch with increasing number of parties, data samples, features, and filters. The computation is single-threaded in a virtual network with an average network delay of 0.17 ms and 1 Gbps bandwidth on 10 Linux servers with Intel Xeon E5-2680 v3 CPUs running at 2.5 GHz with 24 threads on 12 cores and 256 GB RAM.

When the number of data samples is constant per party, *PriCell*’s computation time remains almost constant and only the communication overhead increases when increasing *N* (Figure 5b).

We further analyze *PriCell*’s scalability for *N* = 10 when varying the number of global samples (*s*), the number of features (*m*), and the number of filters (*h*). In Figure 5c, we show that *PriCell* scales linearly when increasing the number of global samples with *N* = 10. Increasing the number of features and filters has almost no effect on *PriCell*’s runtime, due to our efficient packing strategy that enables SIMD operations through features and filters. However, we note that the increase in *h* = 64 in Figure 5e is due to increasing the cryptosystem parameter *𝒩* to have a sufficient number of slots to still rely on our one-cipher packing strategy. The increase in runtime is still linear with respect to *𝒩* and, as expected, the use of larger ciphertexts also produces a slight increase in the communication.

### Comparison to prior work

The most recent solutions for privacy-preserving federated learning in the N-party setting use differential privacy (DP), secure multiparty computation (SMC), or homomorphic encryption (HE).

DP-based solutions^20–22^ in the medical domain introduce noise in the intermediate values to mitigate adversarial FL attacks. However, it has been shown that training an accurate model with DP requires a high-privacy budget^39^. Thus, DP-based solutions introduce a privacy-accuracy trade-off by perturbing the model parameters, whereas *PriCell* decouples the accuracy from the privacy and achieves privacy-by-design with a reasonable overhead.

SMC-based solutions^24–28^ often require the data providers to communicate their data outside their premises to a limited number of computing nodes, and these solutions assume an honest majority among the computing nodes to protect the data and/or model confidentiality. Comparatively, *PriCell* scales efficiently with *N* parties and permits them to keep their data on their premises, withstanding collusions of up to *N −* 1 parties.

Lastly, HE-based solutions for privacy-preserving analytics in distributed medical settings ^31, 32^ allow for functionalities (e.g., basic statistics, counting, or linear regression) different than those *PriCell* enables, and they do not enable the efficient execution of neural networks in a federated learning setting. Due to the fact that the underlying system, the threat model, and the enabled functionalities of all the aforementioned solutions are different from *PriCell*, a quantitative comparison with these works is a challenging task.

We build on the system and threat model proposed by Sav et al.^33^ for enabling privacy-preserving federated learning for neural networks by relying on MHE and make a quantitative comparison with POSEIDON. *PriCell* improves upon the state-of-the-art solution, POSEIDON, by at least one order of magnitude when training CellCnn with the same number of epochs and filters. This is due to *PriCell*’s design for optimizing the use of *Single Instruction, Multiple Data* (SIMD) operations, with a packing strategy that enables encrypting all samples of a batch in a single ciphertext whereas, POSEIDON packs the samples within a batch in different ciphertexts. For a local batch size of 1, 8 filters, 38 features, and 200 cells per sample, *PriCell*’s local computation time is 1.7s; whereas, POSEIDON’s is 15.4s. Increasing the batch size to 100 results in a 100x slower local execution for POSEIDON, whereas it remains constant for *PriCell*, as all samples are packed in one ciphertext. In summary, increasing the batch size or the number of filters yields a linear increase in the advantage of our solution, in terms of local computation time.

### Downstream Analysis

The training in the original CellCnn study aims at detecting rare disease-associated cell subsets via further analysis^35^. Assuming the end model is decrypted upon pre-agreement to conduct these analyses, we further investigate how the changes that we introduce in the CellCnn architecture (see Local Neural Network Operations in the Methods section) affect the detection capability. To be able to make a comparison with the original study in terms of detection capability, we introduce these changes in the original implementation of CellCnn, simulate our encryption, and evaluate the impact in the subsequent analyses. We report our results and about how our changes to the circuit and training affect the detection capability on rare CMV infection, in the Supplementary Note 6.

## Discussion

In this work, we present *PriCell*, a system that enables privacy-preserving federated neural network learning for healthcare institutions, in the framework of an increasingly relevant single-cell analysis, by relying on multiparty homomorphic encryption (MHE). To the best of our knowledge, *PriCell* is the first solution to enable the training of convolutional neural networks with *N* parties under encryption on single-cell data. Using MHE, our solution enables the parties to keep their data on their local premises and to keep the model and any federated learning intermediate values encrypted end-to-end throughout the training. As such, *PriCell* provides security against inference attacks to federated learning^15–19^. *PriCell* also protects the querier’s (researcher’s) evaluation data by using oblivious prediction. The underlying encryption scheme of *PriCell* provides post-quantum security and does not degrade the utility of the data, contrarily to differential privacy-based solutions^20,40^.

In this work, we demonstrate the flexibility of *PriCell* with the different learning parameters (e.g., batch size, number of features, number of filters), different real-world datasets, and a varying number of parties. Our empirical evaluation shows that *PriCell* is able to efficiently train a collective neural network with a large number of parties while protecting the model and the data through homomorphic encryption. We also show that *PriCell*’s computation and communication overhead remains either constant or scales linearly with the number of parties and with the model parameters.

Furthermore, we show that *PriCell* achieves classification accuracy comparable to the centralized and non-encrypted training. Our evaluation demonstrates a substantial accuracy gain by collaboration between the parties when compared to locally training with their data only.

As data sharing in the healthcare domain is usually prevented due to the sensitive nature of data, and due to privacy regulations such as HIPAA^7^ or GDPR^6^, *PriCell* brings unprecedented value for the healthcare domain, exemplified in this work for single-cell analysis, where the data is scarce and sparse. These benefits are extensible to federated healthcare scenarios that rely on machine learning, and constitutes an important landmark for real-world applications of collaborative training between healthcare institutions while preserving privacy.

## Methods

In this section, we detail *PriCell*’s system and threat model, which is based on the state-of-the-art privacy-preserving framework^33^, we introduce some fundamentals on multiparty homomorphic encryption (MHE), describe the datasets used in our accuracy evaluation, and provide a high-level description of local neural network operations under encryption, as well as our experimental setting. We list the frequently used symbols and notations in Supplementary Note 2.

### System and Threat Model

In *PriCell*’s scenario, there are *N* healthcare institutions (parties), each holding its own patient dataset and collectively training a neural network model, without sharing/transferring their local data. Our aim is to preserve the confidentiality of the local data, the intermediate model updates in the FL setting, the querier’s evaluation data, and optionally the final model (see Figure 1a for the system overview). We rely on a synchronous learning protocol, assuming that all parties are available throughout the training and evaluation executions. We note here that this assumption can be relaxed by using different HE schemes, such as threshold or multi-key HE^41,42^, but with a relaxed security assumption for the former and an increased computation cost for the latter.

We consider an *N ™*1 passive-adversary threat model. Hence, we assume that all parties follow the protocol, and up to *N ™*1 colluding parties must not be able to extract any information about the model or other party’s input data. We rely on multiparty homomorphic encryption (MHE) to meet these confidentiality requirements under the posed threat model. In the following section, we briefly introduce the background on the used MHE scheme.

### Multiparty Homomorphic Encryption (MHE)

We rely on a variant of the Cheon-Kim-Kim-Song (CKKS)^43^ cryptographic scheme that is based on the *ring learning with errors* (RLWE) problem (also post-quantum secure ^44^) and that provides approximate arithmetic over vectors of complex numbers, i.e., C*𝒩 /*2 (a complex vector of *𝒩 /*2 slots). Encrypted operations over C*𝒩 /*2 can be carried out in a Single Instruction Multiple Data (SIMD) fashion, which allows for excellent amortization. Therefore, adopting an efficient packing strategy that maximizes the usage of all the slots has a significant impact on the overall computation time.

Mouchet et al.^45^ show how to construct a threshold variant of RLWE schemes, where the parties have their own secret key and collaborate to establish the collective public keys. In this setting, a model can be collectively trained without having to share the individual secret keys of the parties; this prevents the parties’ decryption functionality, without the collaboration of all the parties.

Note that a fresh CKKS ciphertext permits only a limited number of homomorphic operations to be carried out. To enable further homomorphic operations, a ciphertext must be refreshed with a *bootstrapping* operation, once it is exhausted. This operation is a costly function and requires communication in our system, hence we optimize the circuit to minimize the number of bootstrapping operations and the number of ciphertexts to be bootstrapped (see Detailed Neural Network Circuit under this section).

### Datasets

We detail the features of the three used datasets:

#### Non-inflammatory neurological disease (NIND), relapsing–remitting multiple sclerosis (RRMS)

We rely on a large cohort of peripheral blood mononuclear cells (PBMCs), including 29 healthy donors (HD), 31 NIND, and 31 RRMS donors^38^. The dataset comprises samples with a varying number of cells for each donor and 35 markers for each cell. We use this dataset for two classification tasks: (i) HD vs. NIND, and (ii) HD vs. RRMS, as shown in Figure 3 and 4. For both NIND and RRMS experiments and in all experimental settings, we use 48 donors (24 HD, 24 NIND / RRMS) for training and 12 donors (5 HD, 7 NIND / RRMS) for testing.

#### Cytomegalovirus Infection (CMV) classification

We use a mass cytometry dataset^36^ for the classification of cytomegalovirus infection (CMV). This dataset comprises samples from 20 donors with a varying number of cells for each donor and mass cytometry measurements of 37 markers for each cell and has 11 CMV- and 9 CMV+ labels. We use 14 donors for training and 6 donors as a test set in all experimental settings.

#### Acute Myeloid Leukaemia (AML)

We rely on the mass cytometry dataset from Levine et al.^37^ for the 3-class classification problem for healthy, cytogenetically normal (CN), and core-binding factor translocation (CBF). For each cell, the dataset includes mass cytometry measurements of 16 markers. As in the original work^35^, we use the AML samples with at least 10% CD34+ blast cells with the availability of additional cytogenetic information. The final training dataset comprises 3 healthy bone marrows (BM1, BM2, BM3), 2 CN samples (SJ10,SJ12), and 2 CBF samples (SJ1, SJ2). The test set in all experimental settings comprises 2 healthy bone marrows (BM4, BM5), 1 CN (SJ13), and 3 CBF (SJ3, SJ4, SJ5) samples.

The individual donors in all aforementioned training sets are then evenly distributed among *N* parties for *PriCell* collective training. To construct our baselines and to make a fair comparison with the baseline, we use the same data preprocessing for all experiments per setting (centralized CellCnn, Local, or *PriCell*). We give the details of the data preprocessing and parameter selection, in Supplementary Note 3.

### Local Neural Network Operations

In this section, we give a high-level description of the neural network circuit that is evaluated in the encrypted domain (a detailed and step-by-step description can be found in the Supplementary Note 4). We first present the changes introduced to the original CellCnn circuit to enable an efficient evaluation under encryption: (i) we approximate the non-polynomial activation functions by polynomials by using least-squares approximation, (ii) we replace the max pooling with an average pooling, (iii) we replace the ADAM optimizer with the stochastic gradient descent (SGD) optimizer with mean-squared error and momentum acceleration. Finally, we introduce the packing strategy used in *PriCell* and give a high-level circuit overview. We give more details on these steps and optimizations, in the Supplementary Note 4, and we empirically evaluate the effect of these optimizations on the model accuracy in a distributed setting in the Results section.

#### Polynomial Approximations

With additions and multiplications, the CKKS scheme can efficiently evaluate polynomials in the encrypted domain. However, these two basic operations are not sufficient for easily evaluating non-linear functions, such as sign or sigmoid. A common strategy to circumvent this problem is to find a polynomial approximation of the desired function. We rely on polynomial least-squares approximations for the non-polynomial activation functions, such as sigmoid, and we use identity function after convolution (instead of ReLU). We show in our Results section that these changes only have a negligible effect on the model accuracy.

#### Pooling

The original CellCnn circuit makes use of both max pooling and average pooling. Max pooling requires the computation of the non-linear sign function which cannot be efficiently done under encryption. We replace the max pooling with the average pooling, which is a linear transformation and brings the following advantages: (i) it is efficient for computing under encryption with only additions and constant multiplication, (ii) it simplifies the backward pass under encryption, (iii) it commutes with other linear transformations or functions, such as the convolution and the identity activation, which allows for an efficient preprocessing of the data and reduces the online execution cost. Indeed, we are able to pre-compute the average pooling on the data, which reduces the input size of a batch of samples from *n ×c ×m* to *n ×m*, i.e., we remove the dependency on *c*.

#### Optimizer

The original CellCnn training relies on the ADAM optimizer, which requires the computation of square roots and inverses. Although approximating these operations is possible, a high-precision approximation requires an excessive use of ciphertext levels and significantly reduces the efficiency of the training. To avoid these costly operations, we rely instead on the SGD optimizer with momentum acceleration that, for an equivalent amount of epochs, shows a comparable rate of convergence to the ADAM optimizer.

#### Packing Strategy

The CKKS scheme provides complex arithmetic on C*𝒩 /*2 in a SIMD fashion. The native operations are addition, multiplication by a constant, multiplication by a plaintext, multiplication by a ciphertext, *slots rotation* (shifting the values in the vector), and complex conjugation. As the rotations are expensive, when considering encrypted matrix operations, one of the main challenges is to minimize the number of rotations, which can be done by adopting efficient packing strategies and algorithms. We give more details about the packing and algorithms, in the Supplementary Note 4.

With the aforementioned pre-computed pooling, only a 1 *×m* vector is needed to represent a sample, instead of a *c ×m* matrix. Hence, we pack an entire batch of *n* samples in a single ciphertext and compute the forward and backward pass on the whole batch in parallel, which reduces the complexity of the training proportionally to the size of the batch.

#### Encrypted Circuit Overview

Given a batch size of *n* samples, each sample being a matrix *L*_*c×m*_, for *c* the number of cells per sample and *m* number of features (markers) per cell, we first evaluate the mean pooling across the cells in plaintext. The result is a set of *n* vectors of size 1 *×m*, which is packed in an *L*_*n×m*_ matrix. The 1D convolution is evaluated with a *L*_*n×m*_ *×C*_*m×k*_ matrix multiplication. We feed the result to the dense layer *W*_*k×o*_ where *o* is the number of output labels. Lastly, we perform an approximated activation function to the output of the dense layer. Our encrypted circuit with reduced complexity is

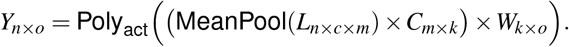

### Experimental Settings

We implemented our solution in Go^46^ by using the open-source lattice-based cryptography Lattigo^47^. We use the implementation of CellCnn^35^ to preprocess the data and to construct baselines. We use Onet^48^ to build a decentralized system and Mininet^49^ to evaluate our system in a virtual network with an average network delay of 0.17 ms and 1 Gbps bandwidth on 10 Linux servers with Intel Xeon E5-2680 v3 CPUs running at 2.5 GHz with 24 threads on 12 cores and 256 GB RAM. The parties communicate over TCP with secure channels (TLS). We choose our security parameters that achieve at least 128-bits security^50^.

## Acknowledgements

We would like to thank Apostolos Pyrgelis, David Froelicher, and Sylvain Chatel who gave valuable feedback on the manuscript. We also thank Shufan Wang and Joao Sa Sousa for their contribution on the experiments and benchmarking. This work was partially supported by the grant #2017-201 of the Strategic Focal Area “Personalized Health and Related Technologies (PHRT)” of the ETH Domain.

## Author Contributions Statement

S.S, J.R.T.-P., M.C, and J.-P.H. conceived the study. S.S. and J.-P.B. developed the methods, implemented them and performed the experiments. All authors contributed to the methodology and wrote the manuscript.

## Data Availability

This study uses previously published data sets^36–38^. We refer the reader to Arvaniti and Claassen^35^ (https://zenodo.org/record/5597098#.YXbaz9ZBzt0) to obtain the CMV and AML datasets, and to Galli et al.^38^ for the repository including the samples from healthy, NIND, and RRMS donors (http://flowrepository.org/experiments/2166/).

## Competing interests

J.R.T.-P. and J.-P.H are co-founders of the start-up Tune Insight SA (https://tuneinsight.com). All authors declare no other competing interests.

## Notes

http://flowrepository.org/experiments/2166/

https://zenodo.org/record/5597098#.YXbaz9ZBzt0

### Competing Interest Statement

Juan R. Troncoso-Pastoriza and Jean-Pierre Hubaux are co-founders of the start-up Tune Insight SA (https://tuneinsight.com). All authors declare no other competing interests.

### Summary of Updates

We have fixed several cross-referencing errors in the main text and updated the author list such that Manfred Claassen and Jean-Pierre Hubaux appear as co-senior authors.

