## Supplementary Material for "Privacy-Preserving Federated Neural Network Learning for Disease-Associated Cell Classification"

### Supplementary Note 1: Cryptography Glossary

Here, we provide a summary of the cryptography terms that are frequently used throughout this paper.
*Multiparty Homomorphic Encryption:* a set of protocols that enable a group of parties to securely compute joint functions over their private inputs by using homomorphic encryption. Compared to LSSS-based (linear secret-sharing scheme) approaches, these protocols scale linearly with the number of parties and do not require private channels.
*Ring Learning With Errors (RLWE):* a computational problem based on the difficulty of solving linear equations that are perturbed by an error. The security of the cryptographic schemes used in this work is based on this problem.
*Ring Degree ($\mathcal{N}$)*: the degree of the RLWE cyclotomic polynomial $X^{\mathcal{N}}+1$.
*Packing:* the act of encrypting multiple scalar values in a single ciphertext by using ciphertext slots.
*Secret Key:* a secret value used to decrypt a ciphertext and to generate the encryption key and evaluation keys.
*Collective Public Key:* a public key generated with the interaction of a set of parties and that can be used by any party to encrypt. The decryption of a ciphertext that is encrypted with the collective key requires all parties to participate in the decryption protocol.
*Evaluation Keys:* special public keys used during the homomorphic evaluation of a circuit (e.g., homomorphic slot rotations).
*Ciphertext Slots:* available space in a ciphertext to encrypt multiple values. In CKKS, the maximum number of *slots* that a ciphertext can have is half of the dimension of the ring degree, i.e. $\mathcal{N}/2$.
*Slots Rotation:* cyclic shift of the values encrypted in a ciphertext.
*Single Instruction, Multiple Data (SIMD) Operations:* the ability to carry out operations in parallel on a batch of data that is encrypted in one ciphertext by using ciphertext slots, in the context of this work.
*Bootstrapping:* the act of homomorphically refreshing a ciphertext to allow for further computations.
*Distributed Bootstrapping:* bootstrapping that requires interaction between the parties but that is less computationally expensive than its non-interactive variant.
*Collective Key Switching:* an interactive re-encryption of a ciphertext to a different secret key.

### Supplementary Note 2: Symbols and Notations

Supplementary Table 1 summarizes the symbols and notation used throughout the paper.

### Supplementary Note 3: Data Preprocessing and Parameter Selection

Our data preprocessing is similar to the one used in CellCnn[35]. To address the distributed setting, we split individual donors in the training set to $N$ institutions. Each party then generates the multi-cell inputs similar to CellCnn (Figure 1b) by selecting $c$ cells per sample, and $z$ samples per class or per patient, depending on the experiment. As a result, each multi-cell input sample has a size of $c \times m$, where $m$ is the number of markers; and the total training set per party has $o \times z$ multi-cell inputs, where $o$ is the number of labels when the data is generated in a per class-basis. The total training set has $p \times z$ multi-cell inputs, where $p$ is the number of patients in that institution when the data is generated in a per patient-basis.

For accuracy evaluation, we use two test datasets for each experimental setting: One is generated by multi-cell inputs of $c$ cells and $z$ samples drawn from test set as in training set, and one is generated with the $g$ of cells per individual to predict the phenotype where $g$ is the minimum of all available cells per individual in the test set. Lastly, for a fair comparison, we use the same test set generated in all settings per dataset.

For all experimental settings, we scale and standardize the marker distributions, based on the training data.

Below, we give the parameters for each experimental setting.

**RRMS/NIND experiments.** For RRMS and NIND experiments in Figures 3 and 4, we generate two training datasets: (i) multi-cell inputs with 100 cells were drawn for each class label to generate a dataset of 30000 samples (phenotype-based) and (ii) multi-cell inputs with 2000 cells were drawn from each patient to generate 480 samples (patient-based). We report the median accuracy in Figures 3 and 4, for patient-based multi-cell input generation in Supplementary Table 2.

The size of the test set for multi-cell inputs, is set to 10000 for the phenotype-based multi-cell generation, and to 96 for patient-based multi-cell generation setting. The test set for phenotype classification is 12 donors for all RRMS and NIND experiments.

**CMV experiments.** We transform the marker measurement with the inverse hyperbolic sine function with a cofactor of 5. The training and test datasets respectively comprise 14 and 6 donors.

For the CellCnn results in Table 2, 200 cells were drawn for each class label to generate one multi-cell sample and 2000 samples are generated per label. For all distributed settings, we also generate 200 cells per sample and gradually decrease the number of samples bagged in each party for a fair comparison.

The size of the test set including multi-cell inputs is set to 4000 and the size of the test set for phenotype classification is 6 donors.

**AML Experiments.** For the AML experiments in Supplementary Table 2, we draw 200 cells per class label to generate

| Notation | Description |
|---|---|
| $P_i$ | $i^{th}$ Party |
| $X_i$ | Input matrix of $P_i$ |
| $y_i$ | True labels of $P_i$ |
| $N$ | Total number of parties |
| $s$ | Total number of samples over parties |
| $L_{n\times c\times m}$ | Batch of samples |
| $C_{m\times h}$ | Convolution filters (weights) |
| $W_{h\times o}$ | Weight matrix of dense layer |
| $n$ | Number of data samples in a batch (batch size) |
| $c$ | Number of cells in multi-cell input |
| $m$ | Number of features (markers) |
| $h$ | Number of filters |
| $o$ | Number of labels (phenotype) |
| $\eta$ | Learning rate |
| $\mu$ | Momentum |
| $\odot$ | Element-wise multiplication |
| $\times$ | Matrix or vector multiplication |
| $\|\|$ | Concatenation |
| **Cryptographic Parameters and Operations** | |
| $\boldsymbol{A}^{\ell}$ | Encryption of $A$ (bold-face) at level $\ell$ |
| $\mathcal{N}$ | Ring degree |
| $R_Q$ | The ring $\mathbb{Z}_Q[X]/(X^{\mathcal{N}}+1)$, with $\mathcal{N}=2^d$ |
| $\mathsf{MultImag}(\cdot)$ | Multiply the slots by the imaginary unit $i$ |
| $\mathsf{Conjugate}(\cdot)$ | Complex conjugate of the slots |
| $\mathsf{Rotate}_i(\cdot)$ | Rotate the slots by $i$ to the left |
| $\mathsf{InnerSum}_{i,j}(\cdot)$ | Sum $j$ batches of $i$ slots |
| $\mathsf{Replicate}_{i,j}(\cdot)$ | Replicate batches of $i$ slots $j$ times |

**Supplementary Table 1.** Frequently Used Symbols and Notations.

multi-cell samples and 1000 samples generated per label. Note that there are 3 class labels for this set of experiments. For all distributed settings, we generate 200 cells per sample and gradually decrease the number of samples, as in other experimental settings. The size of the test set including multi-cell inputs, is set to 3000 and the size of the test set for phenotype classification is 6 donors.

**Machine Learning Parameters.** For all accuracy experiments, we use 8 filters, average pooling, 1 local iteration per party before aggregating local gradients, identity activation after convolution, and an approximated sigmoid activation for the dense layer. For the baseline (CellCnn), we rely on their original optimizer, ADAM, and for *PriCell*, we use SGD with momentum ($\mu$) to enable efficient training of the neural network under encryption. We vary $\mu = 0.5-0.9$ and the learning rate $\eta = 0.0001-0.01$ for the distributed setting.

For the RRMS and NIND experiments, we use a batch size of 64 for the baseline (CellCnn) and gradually decrease the batch size proportionally to the number of parties when data is distributed. For example, when the number of parties is 4, the local batch size is 16. We use the approximated sigmoid activation in $[-1,1]$ with a polynomial degree of 3 for the dense layer, and 30 epochs.

For the CMV experiments, we use a batch size of 200 for the baseline (CellCnn) and gradually decrease the batch size proportionally to the number of parties when the data is distributed. For example, when the number of parties is 2, the local batch size is 100. We use an approximated sigmoid activation in $[-3,3]$ with a polynomial degree of 3 for the dense layer, and 20 epochs.

For all AML experiments, we use a batch size of 200 for the baseline (CellCnn) and 100 for the local batch size in the distributed setting with 2 parties. We use an approximated sigmoid activation in $[-3,3]$ with a polynomial degree of 3 for the dense layer, and 20 epochs.

Lastly, for the Local training in Figures 2, 3, and 4 or the Local row in Supplementary Table 2, we use the original CellCnn architecture, with the same baseline parameters and average the accuracy, precision, and recall over $N$ local parties' models.

**Security Parameters.** Unless otherwise stated, all experiments use a cyclotomic polynomial ring of dimension $\mathcal{N} = 2^{15}$ and an initial level $L = 10$, which provides $2^{14}$ slots per ciphertext and allows for a depth-10 circuit before bootstrapping is needed (10 operations to be carried out before bootstrapping). For the inference times given in Table 1, we start with an initial level of 4 as we do not need the backpropagation. This enables a more efficient forward pass as operations carried out on a ciphertext with a lower level are less expensive. All our cryptographic parameters ensure at least 128-bit security level during the training and up to 256-bit security during the inference.

### Supplementary Note 4: Detailed Neural Network Circuit

**Notations.** We denote a batch of samples, the convolution layer and the dense layer matrices by $L_{n\times c\times m}$, $\boldsymbol{C}_{m\times h}$ and $\boldsymbol{W}_{h\times o}$, respectively, with $n$ the number of samples, $c$ the number of cells per sample per batch, $m$ the number of features (markers), $h$ the number of filters and $o$ the number of output classes (labels). When there is no ambiguity, we eliminate the sub-index of the matrices, e.g., $\boldsymbol{C}_{m\times h}$ is often referred to as $\boldsymbol{C}$. We recall that encrypted matrices are denoted in boldface. We denote the plaintext of binary values as *mask*. When multiplied with a ciphertext, *mask* selects specific slots of the ciphertext by setting the other slots to zero. The terms *row*, *column* and *diagonal* encoding of a matrix denote the mapping of a 2D matrix on a 1D vector by concatenating each row, each column or each diagonal of the matrix respectively.

**Convolution With Pre-Pooling.** Given a hyper-cube batch of samples $L_{n\times c\times m}$, we first preprocess $L$ by applying the average pooling across the cells. As the convolution, average pooling and the activation of this step are all linear transformations, their order is interchangeable. This preprocessing reduces the size of the hyper-cube from $n\times c\times m$ to only $n\times m$, thus removing its dependency on $c$. The convolution is computed with a single matrix multiplication $\boldsymbol{P}_{n\times h} = L_{n\times m}\times\boldsymbol{C}_{m\times h}$, with a row-encoded $\boldsymbol{P}_{n\times h}$ matrix where each row stores the result of the convolution layer for one sample. In the rest of this section, we describe how we pack $L_{n\times m}$ and $\boldsymbol{C}_{m\times h}$ in order to enable an efficient convolution through SIMD operations.

We evaluate the convolution with a diagonally-encoded plaintext and row-encoded ciphertext matrix multiplication. As we operate with non-square matrices, we pad the matrix $\boldsymbol{C}$ with the copies of itself until its number of rows reaches $n$. As such, the result will yield $n$ rows, each of $h$ values. With this approach, the convolution can be evaluated with only $m$ plaintext-ciphertext multiplications and additions, and $m-1$ rotations. If $m\times n$ is not a power of two, cyclic rotations of the ciphertext slots will not result in a cyclic rotation of the flattened matrix. Instead, it requires using the *masking* and rotations, which consumes a level. To overcome this, we pad the flattened matrix with additional copies of itself until it reaches a total of $n+(m-1)$ rows (hence the final size of the flattened matrix is $h(n+m-1)$). This enables us, at the expense of more slots used, to simulate a cyclic rotation. Note that those extra rows are removed by the plaintext multiplication by $L$ that also acts as a masking.

We further reduce the number of operations by making use of complex arithmetic, which is natively provided by the CKKS scheme. Using the following, we compute the dot product of $\langle (a_0,a_1),(b_0,b_1)\rangle$ in a single multiplication:

$$(a_0 - ia_1)\cdot(b_0 + ib_1) = (a_0b_0 + a_1b_1) + i(a_0b_1 - a_1b_0).$$

Hence, the convolution of two half-sized complex matrices $L'_{n,m/2}\times B'_{m/2,h}$ is sufficient to compute the convolution of the real matrices $L_{n\times m}\times\boldsymbol{C}_{m\times h}$:

$$\begin{pmatrix} a_{1,1} & \dots & a_{1,m} \\ \vdots & \ddots & \vdots \\ a_{n,1} & \dots & a_{n,m} \end{pmatrix} \times \begin{pmatrix} b_{1,1} & \dots & b_{1,h} \\ \vdots & \ddots & \vdots \\ b_{m,1} & \dots & b_{m,h} \end{pmatrix} \rightarrow \begin{pmatrix} a_{1,1} - ia_{1,2} & \dots & a_{1,m-1} - ia_{1,m} \\ \vdots & \ddots & \vdots \\ a_{n,1} - ia_{n,2} & \dots & a_{n,m-1} - ia_{n,m} \end{pmatrix} \times \begin{pmatrix} b_{1,1} + ib_{2,1} & \dots & b_{1,m} + ib_{2,m} \\ \vdots & \ddots & \vdots \\ b_{n-1,1} + ib_{n,1} & \dots & b_{n-1,m-1} + ib_{n,m} \end{pmatrix}.$$

The extraction of the real part can be done with complex conjugation and addition. The mapping from $\boldsymbol{C}_{m,h}$ to $\boldsymbol{C}'_{m/2,h}$ is straightforward and can be homomorphically computed with $\boldsymbol{C}' = \boldsymbol{C} + \mathsf{Rotate}_h(\mathsf{MultImag}(\boldsymbol{C}))$. Note that it requires $\boldsymbol{C}$ to be padded with an additional row. The encoding of the plaintext matrix $L_{n,m}$ is done by encoding each diagonal of $L'_{n,m/2}$ in a separate plaintext.

The matrix multiplication $L' \times \boldsymbol{C}'$ is then done with

$$\boldsymbol{P}'_{n\times h} = \sum_{i=0}^{m/2-1} L'_{n\times m/2} \odot \mathsf{Rotate}_{2hi}(\boldsymbol{C}'_{m/2\times h}).$$

The result is a row-encoded $n \times h$ complex matrix. We remove its imaginary part $\boldsymbol{P} = \frac{1}{2}(\boldsymbol{P'} + \mathsf{Conjugate}(\boldsymbol{P'}))$ with the $\frac{1}{2}$ factor being pre-applied to $L'$. Because the number of rotations is reduced by a factor of two, the number of rows for the padding must also be readjusted:

$$\underbrace{n}_{\text{result}} + \underbrace{(\lceil m/2 \rceil - 1)\cdot 2}_{\text{rotations}} + \underbrace{1}_{i \text{ repacking}},$$

and the total number of slots used to encode $\boldsymbol{C}_{m\times h}$ is $nh + (\lceil m/2 \rceil - 1)2h + h$. We give an overview of how $\boldsymbol{C}_{n\times h}$ and $\boldsymbol{P}_{n\times h}$ are each encoded on a vector:

$$\boldsymbol{C}_{m\times h} = (\underbrace{\boldsymbol{C}_{(1,1)},\ldots,\boldsymbol{C}_{(1,h)},\boldsymbol{C}_{(2,1)},\ldots,\boldsymbol{C}_{(2,h)},\ldots,\boldsymbol{C}_{m,1},\ldots,\boldsymbol{C}_{(m,h)},\boldsymbol{C}_{(1,1)},\ldots,}_{nh+(m/2-1)2h+h}0,\ldots,0),$$

$$\boldsymbol{P}_{n\times h} = \big(\boldsymbol{P}_{(1,1)},\ldots,\boldsymbol{P}_{(1,h)},\ldots,\boldsymbol{P}_{(n,1)},\ldots,\boldsymbol{P}_{(n,h)},0,\ldots,0\big).$$

**Dense Layer.** The input to the dense layer is a row-encoded $\boldsymbol{P}_{n\times h}$ matrix that is multiplied with the $\boldsymbol{W}_{h\times o}$ matrix. As the matrix $\boldsymbol{P}_{n\times h}$ is row-encoded and requires a homomorphic extraction of its diagonals, the technique used in the convolution step becomes costly for the dense layer. Instead, we use the multiply-then-inner-sum approach, as in POSEIDON[33]. The values of $\boldsymbol{W}_{h\times o}$ are grouped by samples. We first preprocess $\boldsymbol{P}$ by duplicating it $o$ times for each label. This duplication is done with $\log_2(o) + \mathsf{hw}(o) - 1$ rotations. The matrix $\boldsymbol{W}_{h\times o}$ is column-encoded (row-encoding of its transpose), with each of its columns replicated $n$ times (for each sample):

$$\boldsymbol{W}_{h\times o} = \big(\underbrace{(\boldsymbol{W}_{(1,1)},\ldots,\boldsymbol{W}_{(h,1)}),\ldots,(\boldsymbol{W}_{(1,1)},\ldots,\boldsymbol{W}_{(h,1)})}_{n\times h},\ldots,(\boldsymbol{W}_{(1,o)},\ldots,\boldsymbol{W}_{(h,o)}),\ldots,(\boldsymbol{W}_{(1,o)},\ldots,\boldsymbol{W}_{(h,o)}),0,\ldots,0\big).$$

The multiplication $\boldsymbol{U}_{n\times o} = \boldsymbol{P}_{n\times h} \times \boldsymbol{W}_{h\times o}$ is carried on with a single ciphertext-ciphertext multiplication, followed by an inner-sum of batch $n$ and $h$ ($\log_2(h) + \mathsf{hw}(h) - 1$ rotations). The resulting vector has a size of $onh$:

$$\boldsymbol{U}_{n\times o} = \big(\underbrace{(\boldsymbol{U}_{(1,1)},\times,\ldots,\times),\ldots,(\boldsymbol{U}_{(n,1)},\times,\ldots,\times)}_{n\times h},\ldots,(\boldsymbol{U}_{(1,o)},\times,\ldots,\times),\ldots,(\boldsymbol{U}_{(n,o)},\times,\ldots,\times),0,\ldots,0,\underbrace{(\times,\ldots,\times)}_{h-1}\big),$$

with $\times$ denoting unusable by-product values in the ciphertext slots.

**Repacking for Bootstrapping.** We repack the following elements in a single ciphertext for the optimized bootstrapping:

- $\boldsymbol{U}_{n\times o}$: the result of the dense layer, which uses $onh + h - 1$ slots.
- $\boldsymbol{P}_{n\times h}$: the result of the convolution layer, which uses $nh$ slots.
- $\boldsymbol{W}_{h\times o}$: the dense layer matrix, which uses $onh$ slots.
- $\nabla\boldsymbol{W}^{\mathsf{prev}}_{h\times o}$: the updated dense layer weights of the previous backward pass, which uses $onh$ slots.
- $\nabla\boldsymbol{C}^{\mathsf{prev}}_{m\times h}$: the updated convolution layer weights of the previous backward pass, which uses size $nh + (\lceil m/2 \rceil - 1)\cdot 2h + h$ slots.

The repacking is done solely with additions and rotations, concatenating the empty slots of $\boldsymbol{U}_{n\times o}$:

$$\boldsymbol{D}_{\mathsf{repack}} = \boldsymbol{U}_{n\times o} + \mathsf{Rotate}_{-onh}(\boldsymbol{P}_{n\times h}) + \mathsf{Rotate}_{-2onh}(\boldsymbol{W}_{h\times o}) + \mathsf{Rotate}_{-3onh}(\nabla\boldsymbol{W}^{\mathsf{prev}}_{h\times o}) + \mathsf{Rotate}_{-4onh}(\nabla\boldsymbol{C}^{\mathsf{prev}}_{m\times h}).$$

**Bootstrapping and Repacking for Backward Pass.** The goal of this step is to refresh the ciphertext $\boldsymbol{D}_{\mathsf{repack}}$ to a higher level, to enable more computation and to re-arrange its slots optimally for the backward pass.

- $\boldsymbol{U}_{n\times o}$ : We re-order the slots to arrange them first by samples then by classes, and we duplicate each value $h$ times (replacing the non-zero by-product slots):

$$\boldsymbol{U}_{\mathsf{backW}} = \big(\underbrace{(\boldsymbol{U}_{(0,0)},\ldots,\boldsymbol{U}_{(0,0)},\boldsymbol{U}_{(0,1)},\ldots,\boldsymbol{U}_{(0,1)})}_{2h},\ldots,(\boldsymbol{U}_{(n-1,0)},\ldots,\boldsymbol{U}_{(n-1,0)},\boldsymbol{U}_{(n-1,1)},\ldots,\boldsymbol{U}_{(n-1,1)})\big).$$

We note that the size of this vector remains $onh$. $\boldsymbol{U}_{\mathsf{backW}}$ will be used to compute the dense layer error for the updated dense layer weights. We pack an additional copy of $\boldsymbol{U}$, $\boldsymbol{U}_{\mathsf{backC}}$, which is pre-formatted for the convolution layer error and clustered by sample. By computing twice the same values in parallel, but packed in two different ways (one for the dense layer, one for the convolution layer), we avoid expensive and level-consuming repacking procedures, at the cost of more slot usage. Hence for each label, we pad the $nh$ values with $(m/2-1)2h+h$ additional copies of the relevant rows. The used size is therefore $(nh+(m/2-1)2h+h)o$.

$$\boldsymbol{U}_{\mathsf{backC}} = \Big(\underbrace{(\underbrace{\boldsymbol{U}_{(1,1)},\ldots,\boldsymbol{U}_{(1,1)}}_{h},\ldots,\boldsymbol{U}_{(n,1)},\ldots,\boldsymbol{U}_{(n-1,0)},\boldsymbol{U}_{(1,1),\ldots}}_{nh+(m/2-1)2h+h}),\ldots,\big(\boldsymbol{U}_{(1,o)},\ldots,\boldsymbol{U}_{(1,o)},\ldots,(\boldsymbol{U}_{(n,o)},\ldots,\boldsymbol{U}_{(n,o)},\boldsymbol{U}_{(1,o)},\ldots)\big)\Big).$$

- $\boldsymbol{P}_{n\times h}$: The result of the convolution layer, which is an $n\times h$ row-encoded matrix, is re-arranged by duplicating each of its rows for each class of the dense layer (2 in this example) and multiplied by the learning rate ($\eta$).

$$\boldsymbol{P}_{\mathsf{back}} = \eta\,\big(\underbrace{(\boldsymbol{P}_{(1,1)},\ldots,\boldsymbol{P}_{(1,h)},\boldsymbol{P}_{(1,1)},\ldots,\boldsymbol{P}_{(1,h)})}_{2h},\ldots,(\boldsymbol{P}_{(n,1)},\ldots,\boldsymbol{P}_{(n,h)},\boldsymbol{P}_{(n,1)},\ldots,\boldsymbol{P}_{(n,h)})\big).$$

- $\boldsymbol{W}_{h\times o}$: The dense layer matrix, which is an $h\times o$ column-encoded matrix, is re-arranged by padding each column with itself such that each column has a size of $nh+(m/2-1)2h+h$, for a total size of $o(n\times h+(m/2-1)2h+h)$ and is multiplied by the learning rate ($\eta$).

$$\boldsymbol{W}_{\mathsf{back}} = \eta\,\big(\underbrace{(\boldsymbol{W}_{(1,1)},\ldots,\boldsymbol{W}_{(h,1)},\boldsymbol{W}_{(1,1)},\ldots)}_{nh+(m/2-1)2h+h},\ldots,(\boldsymbol{W}_{(1,o)},\ldots,\boldsymbol{W}_{(h,o)},\boldsymbol{W}_{(1,o)},\ldots)\big).$$

- $\boldsymbol{\nabla W}^{\mathsf{prev}}_{h\times o}$: The previous dense layer updated weights, of size $nho$. The format is preserved (column encoded matrix of size $ho$ with each column replicated $n$ times), but the values are multiplied by the momentum ($\mu$).

$$\nabla\boldsymbol{W}_{\mathsf{back}} = \mu\,\big(\underbrace{(\nabla\boldsymbol{W}_{(1,1)},\ldots,\nabla\boldsymbol{W}_{(h,1)}),\ldots,(\nabla\boldsymbol{W}_{(1,1)},\ldots,\nabla\boldsymbol{W}_{(h,1)})}_{n\times h},\ldots,(\nabla\boldsymbol{W}_{(1,o)},\ldots,\nabla\boldsymbol{W}_{(h,o)}),\ldots,(\nabla\boldsymbol{W}_{(1,o)},\ldots,\nabla\boldsymbol{W}_{(h,o)})\big).$$

- $\boldsymbol{\nabla C}^{\mathsf{prev}}_{m\times h}$: The previous convolution layer updated weights, of size $n\times h+(m/2-1)2h+h$. The format is preserved (row encoded matrix, padded), but the values are multiplied by the momentum ($\mu$).

$$\boldsymbol{\nabla C}_{\mathsf{back}} = \mu\,(\underbrace{\boldsymbol{\nabla C}_{(1,1)},\boldsymbol{\nabla C}_{(1,2)},\ldots,\boldsymbol{\nabla C}_{(1,h)},\boldsymbol{\nabla C}_{(2,1)},\ldots,\boldsymbol{\nabla C}_{(2,h)},\ldots,\boldsymbol{\nabla C}_{(m,h)},\boldsymbol{\nabla C}_{(1,1)},\ldots}_{nh+(m/2-1)2h+h}).$$

In summary, the bootstrapped ciphertext contains the following elements:

$$\boldsymbol{D}_{\mathsf{boot}} = \underbrace{U_{\mathsf{backW}}}_{onh}\ \ ||\ \ \underbrace{U_{\mathsf{backC}}}_{o(nh+(m/2-1)2h+h)}\ \ ||\ \ \underbrace{P_{\mathsf{back}}}_{onh}\ \ ||\ \ \underbrace{W_{\mathsf{back}}}_{o(nh+(m/2-1)2h+h)}\ \ ||\ \ \underbrace{\nabla W_{\mathsf{back}}}_{onh}\ \ ||\ \ \underbrace{\nabla C_{\mathsf{back}}}_{nh+(m/2-1)2h+h}\ ,$$

and the total number of slots used in the ciphertext must respect

$$3onh + (2o+1)(nh + (m/2-1)2h + h) \leq \mathcal{N}/2$$

for the ring degree $\mathcal{N}$. Therefore, a bootstrapped ciphertext can hold up to $n = \lfloor (N/(2h) - (2o+1)(m-1))/(5o+1) \rfloor$ samples. For example, given $N = 2^{15}$, $m = 38$, $h = 8$ and $o = 2$, the ciphertext holds 169 samples. This number is smaller than the number of samples that can be repacked in a single ciphertext before the bootstrapping, hence it sets an upper bound for the number of samples that can be trained in a single batch.

**Backward Pass.** The backward pass is computed using the ciphertext $\boldsymbol{D}_{\text{boot}}$. The different values contained in $\boldsymbol{D}_{\text{boot}}$ are accessed via rotations and ciphertext duplication. Masking is used only at the very end to minimize the use of levels. We start by computing the error of the dense layer formatted for the dense layer update ($\boldsymbol{E}_1$) and formatted for the convolution layer ($\boldsymbol{E}'_1$) at the same time:

$$(\boldsymbol{E}_1 || \boldsymbol{E}'_1) = \sigma'(\boldsymbol{U}_{\text{backW}} || \boldsymbol{U}_{\text{backC}}) \odot \left(\sigma(\boldsymbol{U}_{\text{backW}} || \boldsymbol{U}_{\text{backC}}) - (Y_{\text{backW}} || Y_{\text{backC}})\right),$$

with $Y_{\text{backW}} || Y_{\text{backC}}$, the plaintext labels, accordingly encoded and formatted. We then compute in parallel the updated weights of each sample of the dense layer and the partial error of the convolution layer by multiplying $\boldsymbol{E}_1 || \boldsymbol{E}'_1$ with $\boldsymbol{P}_{\text{back}} || \boldsymbol{W}_{\text{back}}$. Note that $\boldsymbol{P}_{\text{back}} || \boldsymbol{W}_{\text{back}}$ can be accessed and aligned with a rotation on $\boldsymbol{D}_{\text{boot}}$.

$$\nabla \boldsymbol{W} || \boldsymbol{E}_0 = (\boldsymbol{E}_1 || \boldsymbol{E}'_1) \odot (\boldsymbol{P}_{\text{back}} || \boldsymbol{W}_{\text{back}}).$$

$\nabla \boldsymbol{W}$ is clustered by samples, hence we add a summation across the $n$ samples to obtain the updated dense layer weights of the batch. The output contains only a single copy, column-encoded, of $\nabla \boldsymbol{W}$, and of size $oh$. An additional step first adds, then masks and extracts, each column of the result and replicates them $n$ times to expand its size back to $onh$ and to match the original encoding format of $W$ (this masking also removes all the unwanted by-product values). $\nabla \boldsymbol{W}_{\text{back}}$ is added to the result to get the final updated weights of the dense layer.

We finalize the computation of $E_0$ by a summation across the labels, reducing its size to $nh + (m/2-1)2h + h$). $E_0$ is already formatted to be multiplied with the plaintext transposed sample matrix $\eta \cdot L^T$ (pre-pooled and multiplied by $\eta$). This step is same as the convolution layer matrix multiplication:

$$\nabla \boldsymbol{C} = \eta \cdot L^T \times \boldsymbol{E}_0.$$

The result is of size $nh$, with no by-product garbage slots due to the plaintext multiplication, but it needs to be extended to a size of $nh + (m/2-1)2h + h$ to comply with the formatting of $\boldsymbol{C}$. This is done by replicating the $nh$ slots until it reaches at least this amount of slots and by masking the overflow of slots. Similarly, $\nabla \boldsymbol{C}_{\text{back}}$ is added to $\nabla \boldsymbol{C}$, and the result is stored as the newly updated weights for the next batch of samples.

We summarize the given steps in Algorithm 1. Note that the algorithm describes the local computations. Then, the parties collectively aggregate and update the global model, which includes the additional step of taking the mean of $\nabla \boldsymbol{W}$ and $\nabla \boldsymbol{C}$ across all the parties.

**Algorithm 1:** The *local computation* algorithm for encrypted CellCnn training. The exponent of encrypted values (e.g., $y$ for $\boldsymbol{C}^y$) denotes the current ciphertext level. Encryption, encoding, and detailed steps of the repacking during the bootstrapping are omitted for clarity and are described in the Methods section. The value $\mathsf{csize}$ represents $nh + (\lceil m/2 \rceil - 1)2h + h$. We give the function definitions in Supplementary Table 1.

**Input:** $X$ and $Y$ set of samples and labels, learning rate $\eta$, momentum $\mu$, batch size $n$, number of iterations $d$, number of features $m$, number of filters $h$, number of labels $o$, $maskW$ a masking vector containing ones in the first $onh$ slots, $maskW_i$ a set of $o$ masking vectors containing ones in the slots $inh$ to $(i+1)nh$ slots for $0 < i < o$, and $maskC$ a masking vector containing ones in the first $\mathsf{csize}$ slots.

**Output:** The encrypted weights $\boldsymbol{C}$ and $\boldsymbol{W}$.

$\boldsymbol{C}^4 \leftarrow \mathsf{Init}(m,h), \boldsymbol{W}^3 \leftarrow \mathsf{Init}(h,o)$ `// Initialize convolution and dense weights`<br>
$\nabla W^5_{\mathsf{prev}}, \nabla C^4_{\mathsf{prev}} \leftarrow 0$ `// Initialize previous updated weights`<br>
**for** $i = 0; i < d; i = i+1$ **do**<br>
    <u>Batch Selection</u><br>
    $X_{\mathsf{batch}} \leftarrow \mathsf{Select}_n(X)$ `// Select a batch of random samples`<br>
    $Y_{\mathsf{batch}} \leftarrow \mathsf{Select}_n(Y)$ `// Select the corresponding labels`<br>
    $L_{\mathsf{pool}} \leftarrow \mathsf{Pre\text{-}pooling}(X_{\mathsf{batch}})$ `// Apply the pre-pooling to the batch`<br>
    <u>Forward Pass</u><br>
    $\boldsymbol{C}^4_{\mathsf{tmp}} \leftarrow \boldsymbol{C}^4 + \mathsf{Rotate}_h(\mathsf{MultImag}(\boldsymbol{C}^4))$ `// Preprocessing for complex matrix multiplication`<br>
    $\boldsymbol{P}^3 \leftarrow \sum_{i=0}^{\lceil m/2 \rceil -1} L^{\mathsf{diag}[i]}_{\mathsf{pool}} \odot \mathsf{Rotate}_{2hi}(\boldsymbol{C}^4_{\mathsf{tmp}})$ `// Convolution`<br>
    $\boldsymbol{P}^3 \leftarrow \mathsf{Replicate}_{nh,o}(\boldsymbol{P}^3)$ `// Replicate the result for each label`<br>
    $\boldsymbol{U}^2 \leftarrow \mathsf{InnerSum}_{1,h}(\boldsymbol{P}^3 \odot \boldsymbol{W}^3)$ `// Dense layer`<br>
    <u>Bootstrapping</u><br>
    $\boldsymbol{D}^2_{\mathsf{repack}} = \boldsymbol{U}^2 + \mathsf{Rotate}_{-nho}(\boldsymbol{P}^3) + \mathsf{Rotate}_{-2nho}(\boldsymbol{W}^3) + \mathsf{Rotate}_{-3nho}(\nabla \boldsymbol{W}^5_{\mathsf{prev}}) + \mathsf{Rotate}_{-4nho}(\nabla \boldsymbol{C}^4_{\mathsf{prev}})$ `// Pack all necessary values in a single ciphertext`<br>
    $\boldsymbol{D}^9_{\mathsf{boot}} \leftarrow \mathsf{Bootstrapp}_{\eta,\mu}(\boldsymbol{D}^2_{\mathsf{repack}})$ `// Refresh the ciphertext and formatting for the backward pass`<br>
    <u>Backward Pass</u><br>
    $\boldsymbol{U1}^7 \leftarrow \sigma(\boldsymbol{D}^9_{\mathsf{boot}})$ `// Activation`<br>
    $\boldsymbol{U2}^7 \leftarrow \sigma'(\boldsymbol{D}^9_{\mathsf{boot}})$ `// Activation derivative`<br>
    $\boldsymbol{E1}^6 \leftarrow \boldsymbol{U2}^7 \odot (\boldsymbol{U1}^7 - Y_{\mathsf{batch}})$ `// Dense layer error`<br>
    $\boldsymbol{P}^9 \leftarrow \mathsf{Rotate}_{nho+o\cdot\mathsf{csize}}(\boldsymbol{D}^9_{\mathsf{boot}})$ `// Access pooling result and dense layer weights`<br>
    $\nabla \boldsymbol{W}^5 \leftarrow \boldsymbol{P}^9 \odot \boldsymbol{E1}^6$ `// Dense layer updated weights and convolution layer error`<br>
    $\boldsymbol{E0}^5 \leftarrow \mathsf{Rotate}_{nho}(\nabla \boldsymbol{W}^5)$ `// Access convolution layer error`<br>
    $\nabla \boldsymbol{W}^5 \leftarrow \mathsf{InnerSum}_{oh,n}(\nabla \boldsymbol{W}^5)$ `// Finish updated weights with summation across the samples`<br>
    $\boldsymbol{E0}^5 \leftarrow \mathsf{InnerSum}_{\mathsf{csize},o}(\boldsymbol{E0}^5)$ `// Finish E1 with summation across the labels`<br>
    $\nabla \boldsymbol{C}^4 \leftarrow \sum_{i=0}^{\lceil n/2 \rceil -1}(0.5 \cdot L^{T,\mathsf{diag}[i]}_{\mathsf{pool}}) \odot \mathsf{Rotate}_{2mi}(\boldsymbol{E0}^5)$ `// Multiply with the transposed samples`<br>
    $\nabla \boldsymbol{C}^4 \leftarrow \nabla \boldsymbol{C}^4 + \mathsf{Conjugate}(\nabla \boldsymbol{C}^4)$ `// Clean imaginary part`<br>
    $\nabla \boldsymbol{C}^4 \leftarrow \mathsf{Replicate}_{mh,\lceil \mathsf{csize}/mh \rceil}(\nabla \boldsymbol{C}^4)$ `// Format updated weights for convolution layer`<br>
    $\nabla \boldsymbol{W}^5 \leftarrow \mathsf{Replicate}_{h,n}(\sum_{i=0}^{o-1} \mathsf{Rotate}_{-inh}(maskW_i \odot \nabla \boldsymbol{W}^5))$ `// Format updated weights for dense layer`<br>
    $\nabla \boldsymbol{W}^8_{\mathsf{prev}} \leftarrow maskW \odot \mathsf{Rotate}_{2nho+2o\cdot\mathsf{csize}}(\boldsymbol{D}^9_{\mathsf{boot}})$ `// Access and extract the previous updated weights`<br>
    $\nabla \boldsymbol{C}^8_{\mathsf{prev}} \leftarrow maskC \odot \mathsf{Rotate}_{3nho+2o\cdot\mathsf{csize}}(\boldsymbol{D}^9_{\mathsf{boot}})$ `// Access and extract the previous updated weights`<br>
    <u>Weights Update</u><br>
    $\nabla \boldsymbol{W}^5 \leftarrow \nabla \boldsymbol{W}^5 + \nabla \boldsymbol{W}^8_{\mathsf{prev}}$ `// Add previous updated weights with momentum`<br>
    $\nabla \boldsymbol{C}^4 \leftarrow \nabla \boldsymbol{C}^4 + \nabla \boldsymbol{C}^8_{\mathsf{prev}}$ `// Add previous updated weights with momentum`<br>
    $\boldsymbol{C}^4 \leftarrow \boldsymbol{C}^4 - \nabla \boldsymbol{C}^4$ `// Update the weights`<br>
    $\boldsymbol{W}^3 \leftarrow \boldsymbol{W}^3 - \nabla \boldsymbol{W}^5$ `// Update the weights`<br>
    $\nabla \boldsymbol{C}^4_{\mathsf{prev}} \leftarrow \nabla \boldsymbol{C}^4$ `// Store the new updated weights`<br>
    $\nabla \boldsymbol{W}^5_{\mathsf{prev}} \leftarrow \nabla \boldsymbol{W}^5$ `// Store the new updated weights`<br>
**end**<br>
**return**

### Supplementary Note 5: Summary of Experiments

In Supplementary Table 2, we show the median accuracy, precision, recall, and F-score values of 10 runs for RRMS and NIND experiments with patient-based sub-sampling shown in Figures 3c, 3d and Figures 4c, 4d, and CMV experiments for phenotype-based sub-sampling shown in Figure 2. We also report these metrics for the AML classification for the centralized and two-party *PriCell* settings.

We note that for 3-class classification, i.e., AML, we rely on macro-averaging on metrics, and we calculate the F-score in Supplementary Table 2 over the averaged precision and recall for the Local-training experimental setting.

Our results show that *PriCell* achieves an accuracy comparable to the centralized and non-private solutions. The accuracy achieved by *PriCell* remains almost the same as the centralized one, and the slight decrease in phenotype classification in NIND classification is due to the limited number of samples in the test set for this task, i.e., there are only 12 patients in the NIND phenotype test set, which results in accuracy decrease of 4% in the median value when the trained model misses only one patient classification.

Lastly, we note that the differences in the precision and recall values are due to the nature of the preprocessing and training mechanism: the random selection of multi-cell inputs generates higher or lower precision and recall values depending on the eventual selection, even in the centralized and no privacy-protection solution.

| Setting/Metrics | Accuracy | Precision | Recall | F-score |
|---|---|---|---|---|
| **RRMS (multi-cell / phenotype classification)** | | | | |
| CellCnn | 0.65 / 0.67 | 0.62 / 0.71 | 0.61 / 0.71 | 0.66 / 0.71 |
| Local ($N$=2) | 0.59 / 0.62 | 0.59 / 0.68 | 0.62 / 0.64 | 0.59 / 0.66 |
| *PriCell* ($N$=2) | 0.65 / 0.67 | 0.66 / 0.75 | 0.62 / 0.64 | 0.64 / 0.67 |
| Local ($N$=4) | 0.55 / 0.55 | 0.57 / 0.61 | 0.60 / 0.61 | 0.58 / 0.62 |
| *PriCell* ($N$=4) | 0.64 / 0.67 | 0.63 / 0.73 | 0.61 / 0.64 | 0.61 / 0.69 |
| Local ($N$=6) | 0.53 / 0.52 | 0.53 / 0.58 | 0.55 / 0.54 | 0.53 / 0.57 |
| *PriCell* ($N$=6) | 0.64 / 0.67 | 0.68 / 0.80 | 0.61 / 0.64 | 0.63 / 0.69 |
| **NIND (multi-cell / phenotype classification)** | | | | |
| CellCnn | 0.72 / 0.75 | 0.82 / 0.83 | 0.75 / 0.71 | 0.76 / 0.77 |
| Local ($N$=2) | 0.57 / 0.58 | 0.65 / 0.65 | 0.59 / 0.52 | 0.62 / 0.63 |
| *PriCell* ($N$=2) | 0.72 / 0.75 | 0.73 / 0.75 | 0.80 / 0.86 | 0.78 / 0.80 |
| Local ($N$=4) | 0.55 / 0.55 | 0.66 / 0.68 | 0.51 / 0.50 | 0.59 / 0.58 |
| *PriCell* ($N$=4) | 0.72 / 0.71 | 0.75 / 0.73 | 0.84 / 0.71 | 0.78 / 0.75 |
| Local ($N$=6) | 0.53 / 0.52 | 0.63 / 0.64 | 0.57 / 0.56 | 0.59 / 0.57 |
| *PriCell* ($N$=6) | 0.71 / 0.71 | 0.73 / 0.75 | 0.76 / 0.71 | 0.75 / 0.75 |
| **CMV (multi-cell / phenotype classification)** | | | | |
| CellCnn | 0.80 / 0.75 | 0.72 / 0.58 | 0.98 / 1.00 | 0.83 / 0.73 |
| Local ($N$=2) | 0.54 / 0.58 | 0.52 / 0.42 | 0.50 / 0.50 | 0.52 / 0.47 |
| *PriCell* ($N$=2) | 0.79 / 0.75 | 0.71 / 0.58 | 0.98 / 1.00 | 0.83 / 0.73 |
| Local ($N$=3) | 0.59 / 0.55 | 0.56 / 0.40 | 0.64 / 0.67 | 0.60 / 0.49 |
| *PriCell* ($N$=3) | 0.79 / 0.75 | 0.76 / 0.58 | 0.84 / 1.00 | 0.80 / 0.73 |
| Local ($N$=5) | 0.50 / 0.52 | 0.44 / 0.31 | 0.57 / 0.55 | 0.50 / 0.39 |
| *PriCell* ($N$=5) | 0.78 / 0.75 | 0.69 / 0.58 | 0.97 / 1.00 | 0.82 / 0.73 |
| **AML (multi-cell / phenotype classification)** | | | | |
| CellCnn | 1.00 / 1.00 | 1.00 / 1.00 | 1.00 / 1.00 | 1.00 / 1.00 |
| Local ($N$=2) | 0.98 / 1.00 | 0.98 / 1.00 | 0.98 / 1.00 | 0.98 / 1.00 |
| *PriCell* ($N$=2) | 1.00 / 1.00 | 1.00 / 1.00 | 1.00 / 1.00 | 1.00 / 1.00 |

**Supplementary Table 2.** Classification performance (accuracy, precision, recall, and F-score) of the models obtained with *PriCell*, original CellCnn, and local training without collaboration for RRMS, NIND, CMV, and AML classification tasks. All models are tested on two datasets for multi-cell and phenotype classification respectively, separated with '/'.

### Supplementary Note 6: Downstream Analysis

The original CellCnn[35] study aims at detecting the rare disease-associated cell subsets via learned filter weights. The final filter weights are used to select phenotype-associated cell subsets via a filter response, i.e., the weighted sum of the abundance profile for each cell. As the cell subset selected by a filter can contain more than one cell type, the authors perform a density-based clustering of the group of cells with high cell-filter responses.

We perform an analogous analysis to evaluate the effect of our introduced changes in the original neural network architecture, namely the average pooling, the approximated activation functions, and the optimizer. We introduce these changes in CellCnn's original implementation, simulate our encryption on centralized data, and conduct further analysis by using their downstream analysis[35]. We use the CMV infection dataset with $c = 200$ cells per multi-cell input and $z = 1000$ samples per phenotype to generate the training dataset. The test set is generated as explained in the Data Preprocessing and Parameter Selection section. We train 20 models for CellCnn and 20 models for *PriCell* simulation and take the best 3 models for each approach based on the validation accuracy, as in the original work[35]. In all model training, we use 20 epochs with early-stopping and varying numbers of filters in each model training.

In Supplementary Figure 1, we show the consensus filters, i.e., one representative filter per class (phenotype) that has minimum distance to all other members of the hierarchically clustered filters, based on a threshold of 0.2, found by CellCnn and *PriCell* simulation, respectively. In both Supplementary Figures 1a and 1b, we observe that the filter which is positively associated (second filter) with previous CMV infection gives more weights to the CD16, CD57, NKG2C, and CD94 markers. We note that while the trend of consensus filters is similar, the distribution of the consensus filters, i.e., the final filter weights and the scale of the values, differs between CellCnn and *PriCell*. This is due to our approximated activation function that affects the final values of the filter weights but does not affect the interpretation from the consensus filters. Similar results were found for the repetition of these experiments, which suggests that, as in the original work[35], our encrypted model is able to find natural killer (NK) cell populations associated with prior CMV infection.

In Supplementary Figure 2, we show the boxplot of the selected cell population frequencies from the test samples of the CMV- and CMV+ classes by using the positively associated filter. Although CellCnn has higher discriminative frequencies, *PriCell* simulation is able to select CMV+ cell populations with the positively associated filter.

Finally, we show in Supplementary Figure 3 the marker expression profiles for all cells vs. cell population selected by the positively associated filter learned by CellCnn training (Supplementary Figure 3a) and by *PriCell* simulation(Supplementary Figure 3b). In both CellCnn and *PriCell* training, we again observe that the positively associated filter weighs CD16, CD57, NKG2C, and CD94 markers more than the others.

In summary, we show that the *PriCell* training does not affect the further findings of an existing work that performs training on a centralized data without integrating a privacy-preserving mechanism.

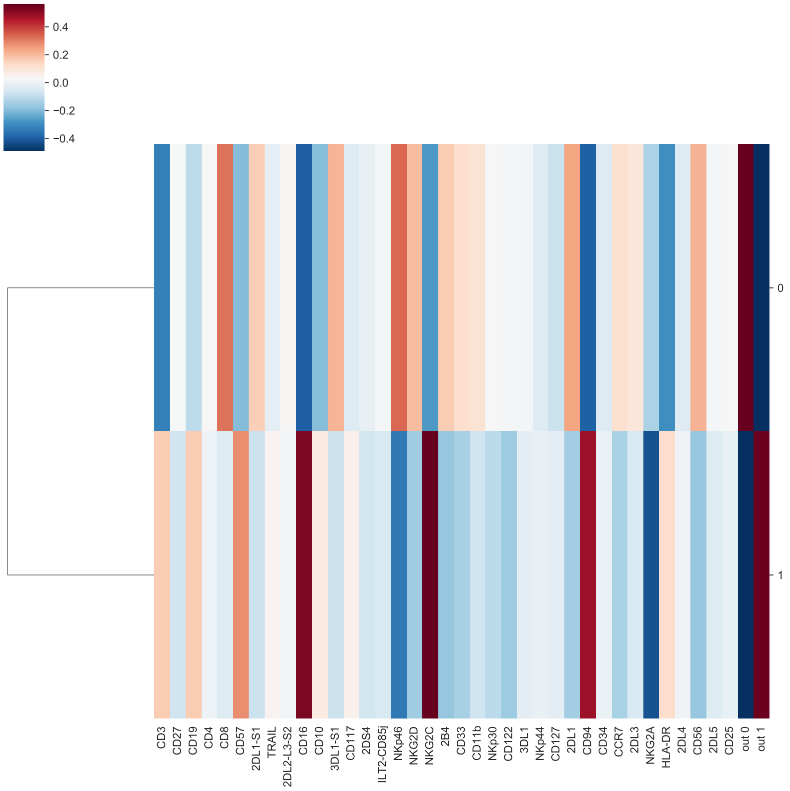


**(a)** Consensus filters found by CellCnn

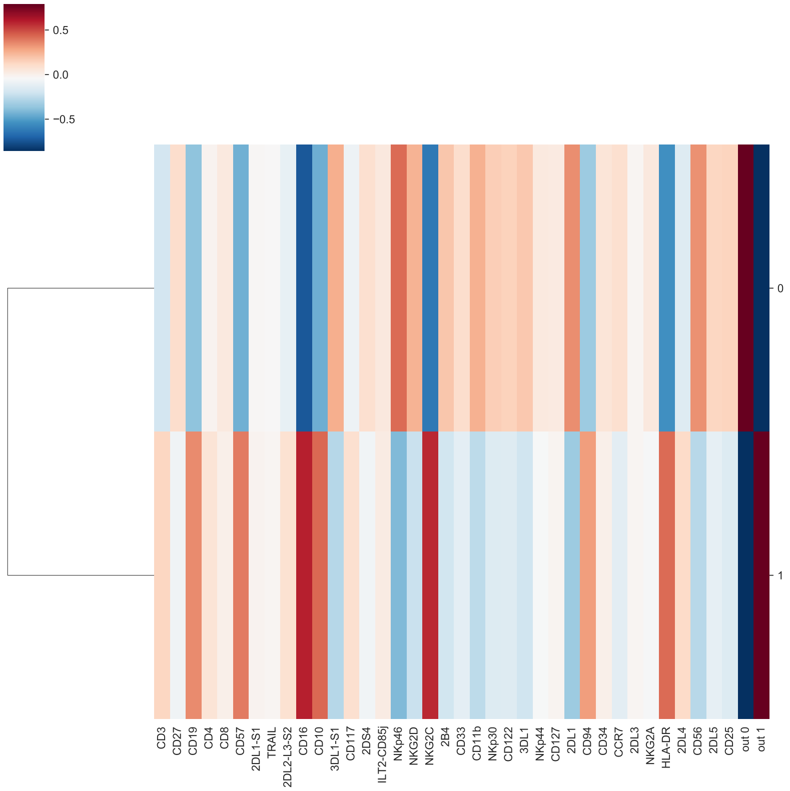


**(b)** Consensus filters found by *PriCell* simulation

**Supplementary Figure 1.** Comparison of the consensus filters (one representative filter per class label) learned by (a) CellCnn original architecture, and by (b) *PriCell*'s adapted architecture for encrypted training on CMV dataset.

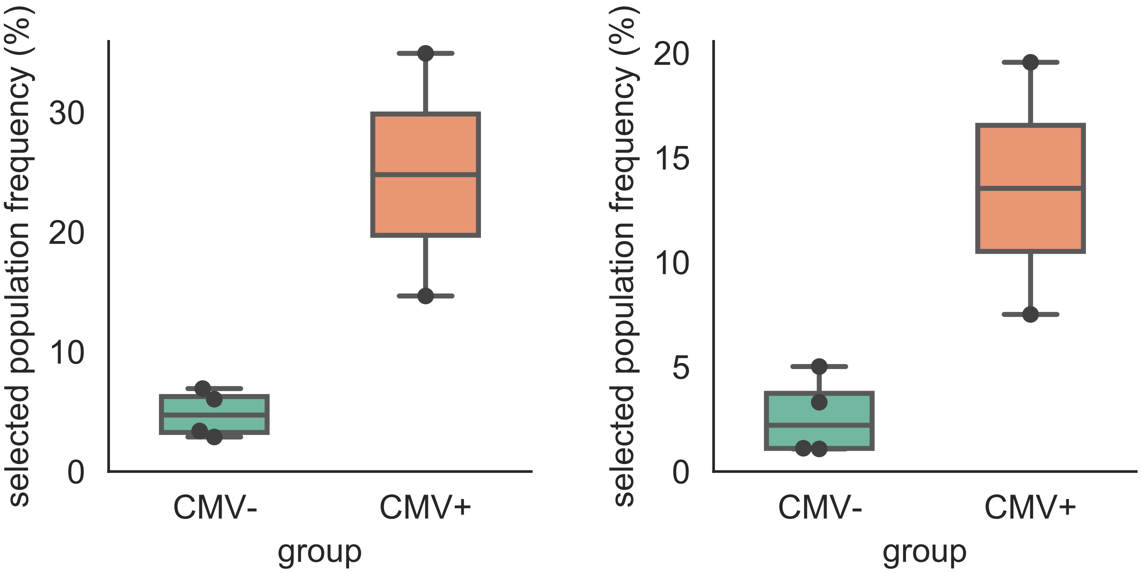


**(a)** CellCnn **(b)** *PriCell* simulation

**Supplementary Figure 2.** Comparison of the selected cell population frequencies from the test samples of the CMV- and CMV+ classes by using the positively associated filter learned by (a) CellCnn original architecture, and by (b) *PriCell*'s adapted architecture for encrypted training.

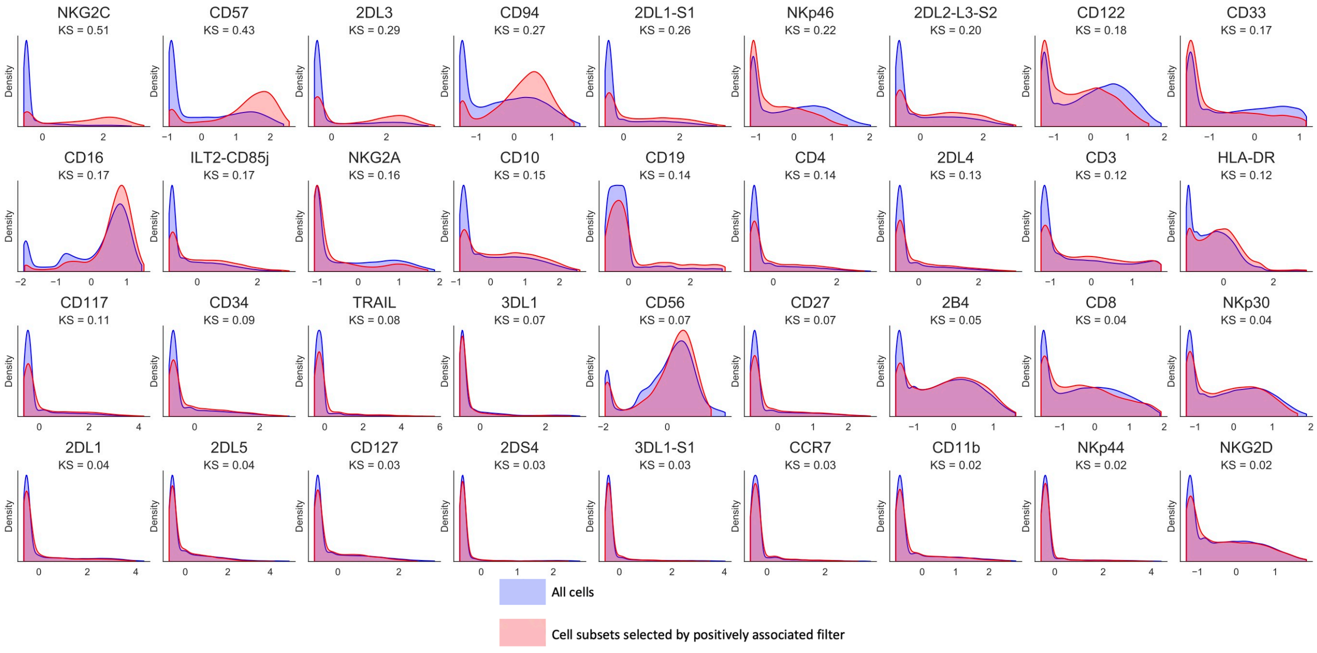


**(a)** CellCnn

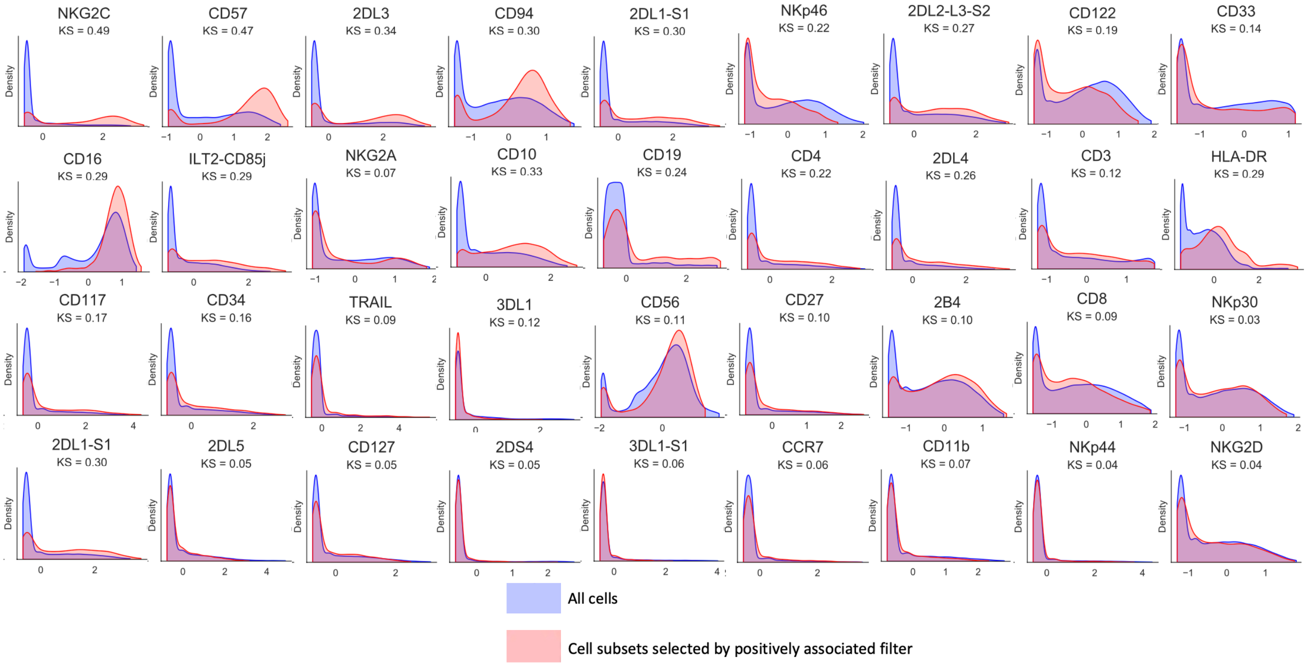


**(b)** *PriCell* simulation

**Supplementary Figure 3.** Comparison of the histograms of univariate z-transformed marker expression profiles for all cells and for the cell population selected by the positively associated filter learned by (a) CellCnn original architecture, and by (b) *PriCell*'s adapted architecture for encrypted training on CMV dataset. The distributions show that *PriCell* training does not affect the findings of the non-privacy preserving training.
